# Tim4 enables large peritoneal macrophages to cross-present tumor antigens at early stages of tumorigenesis

**DOI:** 10.1101/2023.09.14.557776

**Authors:** Sonal Joshi, Lucía López Rodríguez, Luciano Gastón Morosi, Roberto Amadio, Manendra Pachauri, Mauro Giacca, Giulia Maria Piperno, Federica Benvenuti

## Abstract

Cross-presentation of tumor antigens by subsets of macrophages at different stages along tumor progression may have divergent impacts on anti-tumoral immune responses. Here we show that TIM4^+^ large peritoneal macrophages (LPM) avidly capture tumor cells and cross-present tumor-associated antigens at early stages of peritoneal infiltration by ovarian cancer cells. The phosphatidylserine (PS) receptor TIM4 promotes maximal uptake and triggers inflammatory and metabolic gene programs in combination with cytoskeletal remodeling and upregulation of transcriptional signatures related to antigen processing. At the cellular levels, TIM4 is recruited with F-actin at the phagocytic cup and translocates in nascent phagosomes, controlling the kinetic of phagosomal acidification and cargo degradation. TIM4 deletion abrogates cross-presentation of tumor-associated antigens and blunts expansion of effector CD8 T cells at tumor inception. In addition, targeting tumor antigens to LPM by PS liposomes can trigger CD8 T cell activation. Together these results suggest that TIM4 enables LPMs to scan the antigenic content of incoming tumor cells to promote immune surveillance by CD8 T cells and, at defined temporal windows, may be exploited for therapeutic purposes.

## Introduction

Tumor-associated macrophages (TAM) include tissue-resident cells that preexist tumor development and monocyte-derived macrophages that differentiate in situ under tumor-derived cues. While a large body of evidence established the tumor-promoting functions of monocyte-derived TAMs ^1–5^, the precise function of tissue-resident macrophages (TRM) during initiation of primary tumors or early colonization of metastatic sites is still poorly understood. Sparse evidence suggests that, within TRM, subsets with tumor-promoting functions ^6,7^ coexist with subsets endowed with CD8 stimulatory properties ^8,9^. These data indicate that at least a fraction of TRM possesses intrinsic tumor protective functions, depending on tumor tissue and stage, highlighting the importance of elucidating their function and therapeutic potential.

Cross-presentation, the specialized pathway to process and present exogenous tumor antigens on MHC class-I, is particularly efficient in cDC1, a specialized subset invariably linked to anti-tumorigenic properties via activation of anti-tumoral CD8 T cells ^10,11^. Cross-presentation, albeit less efficient on a per-cell basis ^12^, can operate as well in monocyte-derived macrophages ^13,14^ and it is emerging as a critical factor to establish CD8 T cell exhaustion in advanced tumors ^15,16^. Conversely, the role, underlying mechanisms and overall impact of cross-presentation by TRM at initial tumor stages remain poorly explored. The peritoneal cavity, the site of metastasis of gut and ovarian tumors, is dominated by two well-characterized classes of macrophages ^17^. Small peritoneal macrophages (SPM) arise from bone-marrow myeloid precursors, are poorly represented at steady state and expand dramatically during inflammation and tumor progression. Large peritoneal macrophages (LPMs) are the largest population at steady state and derive from embryonic precursors, yet they can be replenished by long-lived bone-marrow cells under Gata6-dependent environmental signals ^18,19^. While LPM intrinsic functions are linked to tissue repair, the subset acquires pro-tumorigenic functions during tumor progression, shifting to high production of reactive oxygen species and autophagic adaptation ^20–22^. LPM, like resident macrophages of the liver, spleen and brain, express high levels of the phosphatidylserine (PS) receptor TIM4, which confers the ability to engulf apoptotic cells, contributing to silent clearance and maintenance of tissue homeostasis ^23,24^. In addition, TIM4 was recently found to regulate cholesterol metabolism in adipose tissue macrophages and during anti-viral responses ^25,26^. In cancer tissues, TIM4 has been associated to disparate, context-dependent functions. Cavity macrophages in advanced metastatic tumors use TIM4 to sequester PS-expressing, activated CD8^+^ T cells, dampening anti-tumoral responses ^27^. Past work suggested that monocytes-derived TAMs infiltrating B16 subcutaneous tumors acquire TIM4 expression, which regulates autophagy and degradation of tumor antigens^28^. Conversely, our group previously uncovered that high expression of TIM4 by lung tissue resident cDC1 in nascent tumors is required to engulf dying cancer cells and to cross-present tumor antigens to initiate anti-tumoral T cell responses ^29^. Human data confirmed that myeloid cells expressing TIM4 in tertiary lymphoid structures positively correlate to better survival across different cancer types ^9,29^. On these bases, we hypothesized that TIM4 highly expressed on LPM may serve to scrutinize antigenic content in incoming metastatic cells. By establishing a metastatic model of ovarian cancer expressing a phagocytic reporter and a model antigen we show that, at initial tumor stages, TIM4 mediates uptake and cross-presentation of tumor antigens, leading to priming of peritoneal CD8^+^ T cells. Mechanistically, we uncover that TIM4 controls the trafficking of ingested dead cells, routing the cargo to cross-presentation competent phagosomes.

## Results

### Uptake of tumor cells is attributed to TIM4-expressing Large Peritoneal Macrophages in peritoneal cavity

To investigate the role of TIM4 on peritoneal resident macrophages, we established a model of ovarian cancer that recapitulates spreading of cancer cells to the peritoneum, based on the syngeneic line ID8 ^30^. ID8 were genetically engineered to express the pH stable reporter ZsGreen and the OVA model antigen (ID8^ZGO^) to allow tracking of phagocytosis and presentation of tumor antigens ^31^. Tumor cells were implanted in the peritoneum and tissues were harvested 24 hr or 15 days after challenge, to examine peritoneal macrophages immediately after seeding and at initial engraftment. Flow cytometry analysis of the peritoneal immune infiltrate showed that LPM and SPM (defined as F4/80^high^, MHCII^low^ and F4/80^low^/MHCII^high^, respectively), expand and accumulate after 15 days, in line with past reports ^22,32^(**Fig 1 A, B**). In addition, we observed a slight induction in the proportion of inflammatory monocytes, neutrophils and CD8^+^ T cells in tumor-challenged mice, paralleled by a reduction in B cells (**Fig S1 A, B**). In line with previous reports, a large fraction of steady state LPM showed strong TIM4 expression, which slightly declined in tumor-challenged animals. Weak TIM4 expression was also present on cDC1, whereas no expression was detected on the other peritoneal phagocytes, including SPM (**Fig 1C**). We next examined acquisition of ZsGreen signal by peritoneal phagocytes as a measure of cancer cells uptake. LPM displayed the highest phagocytic capacity (70.18 %± 9.24) followed by cDC2 (27.87% ± 5.01). In contrast, SPM, Ly6C^+^ monocytes and neutrophils were weakly associated with ZsGreen signal (**Fig 1D, S1C**). Within LPM, the uptake was restricted to the TIM4^+^ fraction both at 24 hr and at 15 days, suggesting a selective ability of this population to engulf tumor cells (**Fig 1E, Fig S1E**). As a control, we verified that cancer cells in the peritoneum express PS, the ligand of TIM4 (**Fig S1D**). Confocal imaging of LPM sorted from ID8-challenged mice confirmed actual internalization of tumor cells by TIM4 positive LPM (**Fig 1F**). Apart from LPM, other subsets of macrophages residing in the peritoneal cavity, such as mesothelium and omentum, have also been shown to play an important role during ovarian cancer progression. Omentum is a fatty tissue that is formed from fold of mesothelium and hosts resident macrophage subset expressing TIM4 ^7^. However, we found TIM4 expression was much weaker on omental CD64^high^ F4/80^high^ resident macrophages than on LPM and further decreased in tumor-challenged animals. In addition, cancer cell uptake was associated to TIM4 negative cells in this context (**Fig S2A-C**). We conclude that LPM expressing high TIM4 levels are especially efficient in capturing cancer cells seeding to the peritoneal cavity.

**Fig 1.**
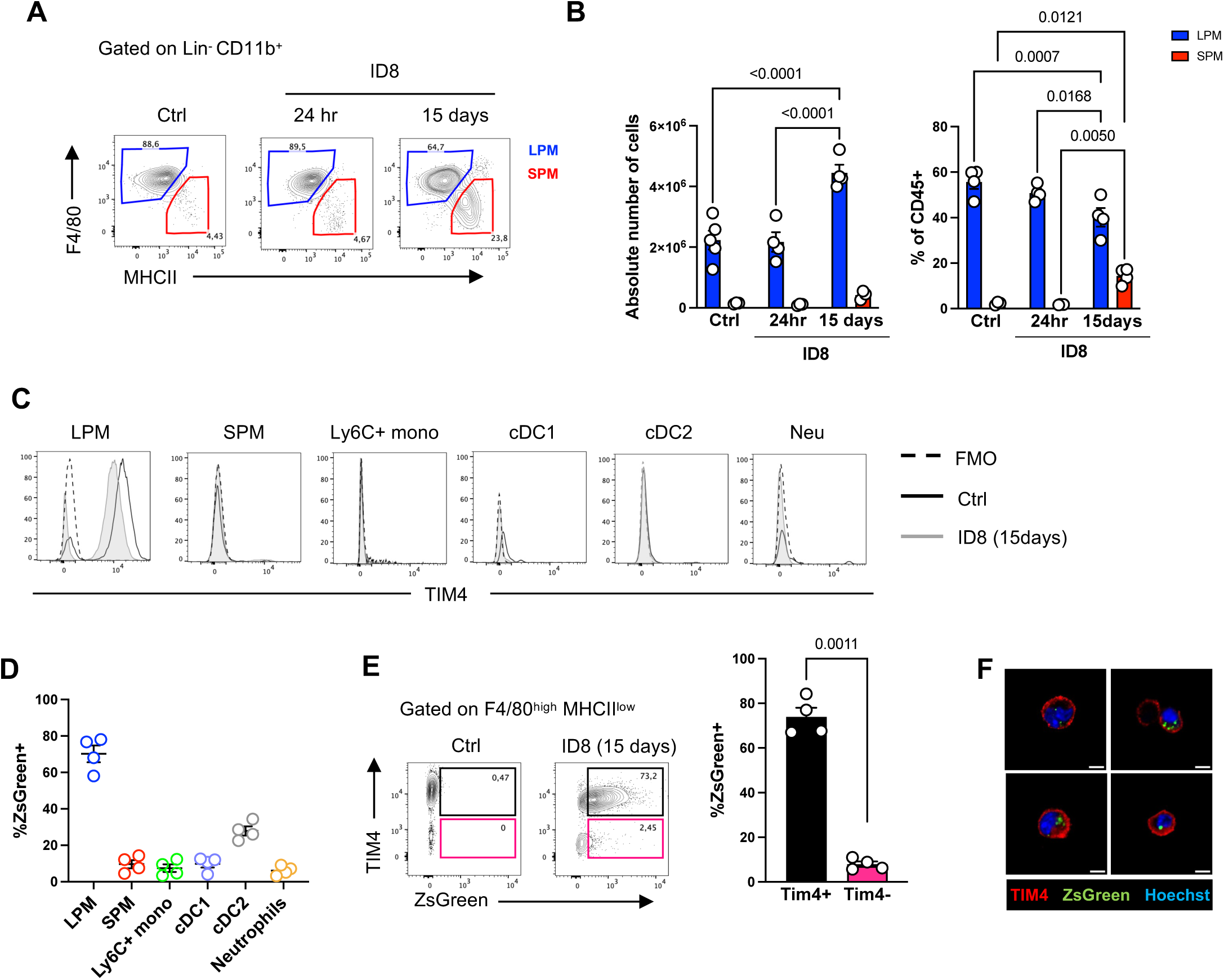
TIM4^high^ LPM efficiently engulf ovarian cancer cells that colonize the peritoneal cavity. **A-B)** Mice were challenged with ID8^ZGO^ and peritoneal cells were collected after 24hr or 15 days. **A)** Representative flow cytometry plots showing LPM (F4/80^high^ MHCIIl^ow^; Blue) and SPM (F4/80^low^ MHCII^high^;Red) macrophages, gated on CD45^+^ CD11b^high^, lineage negative cells (CD19, CD3, B220, NK1.1, Ly6C, Ly6G), in peritoneal cavity of mice 24 hr or 15 days after injection of ID8^ZGO^. **B)** Quantification of (left) absolute numbers and (right) fraction of LPM and SPM in resting (Ctrl) and tumor-bearing (ID8) animals. Data from n=3,4 animals, Data represent mean ± SEM, two-way ANOVA followed by Šídák’s multiple comparisons test. **C)** Representative histograms of TIM4 expression on peritoneal phagocytes at day 15. Large peritoneal macrophages (LPM), small peritoneal macrophages (SPM), Ly6C+ monocytes (Ly6C+ mono), type I dendritic cells (cDC1), type II dendritic cells (cDC2), Neutrophils (Neu). Each peak is overlaid on the corresponding FMO controls (dotted line). **D)** Quantification of tumor cell uptake plotted as % of ZsGreen^+^ cells within each population. n=4, Data represent mean ± SEM from one of two independent experiments. **E)** Representative dot plots and quantifications of uptake at day 15 by TIM4^+^ (black) and TIM4^-^ (pink) LPMs. n=4, data represent mean ± SEM. Statistical analysis was conducted by paired t-test. **F**) Representative confocal images of sorted LPM containing engulfed ZsGreen^+^ cancer cells.

### TIM4 LPM in the peritoneal cavity efficiently cross-present tumor associated antigen

To understand transcriptional programs induced by cancer cell engulfment, we isolated by cell sorting the fraction of LPMs that have taken up ZsGreen^+^ fluorescence, 24 hr after ID8^ZGO^ peritoneal challenge. As a control, we used LPMs from naïve animals, as the ZsGreen^-^ fraction may contain cells that already degraded cargo, confounding the analysis. Bulk RNA seq revealed significant changes and differentially expressed genes induced by cancer cell uptake (**Fig 2A, S3A**, Supplementary Table 1). Gene set enrichment analysis showed prominent enrichment of inflammatory responses in LPM that have engulfed cancer cells, in line with individual DEGs (**Fig 2B and S3A**). Moreover, cancer cell uptake induced metabolic rewiring of LPM, increasing glycolysis and cellular respiration. Interestingly, pathways related to the degradation of ingested antigens and MHC class-I antigen presentation were also enriched in cancer cells loaded LPM (**Fig2B, Supplementary table 2**). Based on these results, we moved to directly test cross-presentation of tumor antigens. Labelling with an antibody specific for the pMHC-I OVA complex showed a clear signal on LPM isolated 24 hr or 15 days after ID8^ZGO^ injection, suggesting that tumor antigens are continuously processed and presented on the cell surface (**Fig2C**). To probe the ability to activate CD8^+^ T cells, we next isolated total LPM, ZsGreen positive and negative LPM and SPM from day 15 ID8^ZGO^ challenged mice and cultured them with OVA-specific CD8^+^ T cells (OT-I). Total LPM and ZsGreen^+^ LPM induced robust T cell proliferation, while the ZsGreen negative fraction failed to induce T cell activation. SPM triggered initial CD8^+^ T cells proliferation, however cells arrested after a few cycles resulting in a low proliferation index (**Fig 2D, E**). Consistently, total LPMs and ZsGreen positive LPMs triggered robust IFNγ secretion by CD8^+^ T cells, whilst SPM and the ZsGreen negative LPM fraction were poorly stimulatory (**Fig 2F**). Of note, at late tumor stages (60 days after challenge), LPMs were significantly less efficient at inducing CD8 T cell activation, indicating that tumor-derived factors modify the inherent properties of peritoneal resident macrophages and blunt the ability to cross-present (**Fig S3B**). To explore whether cross-presentation is a specific property of TIM4^+^ LPM, we next tested lung resident alveolar macrophages (AM), that lack TIM4 expression. LPM and AM equally engulfed ID8^ZGO^ cells (**FigS3C**), however, AMs were largely less efficient than LPMs at inducing the proliferation of OVA-specific CD8 T cells (**Fig 2G**). Together, these results indicate that in the peritoneum TIM4^+^ LPMs possess a superior capacity to acquire and cross-present engulfed tumor antigens. Moreover, AM that lack TIM4 are unable to efficiently cross-present ingested tumor antigens, suggesting a specific role for this receptor in mediating cross-presentation.

**Fig 2.**
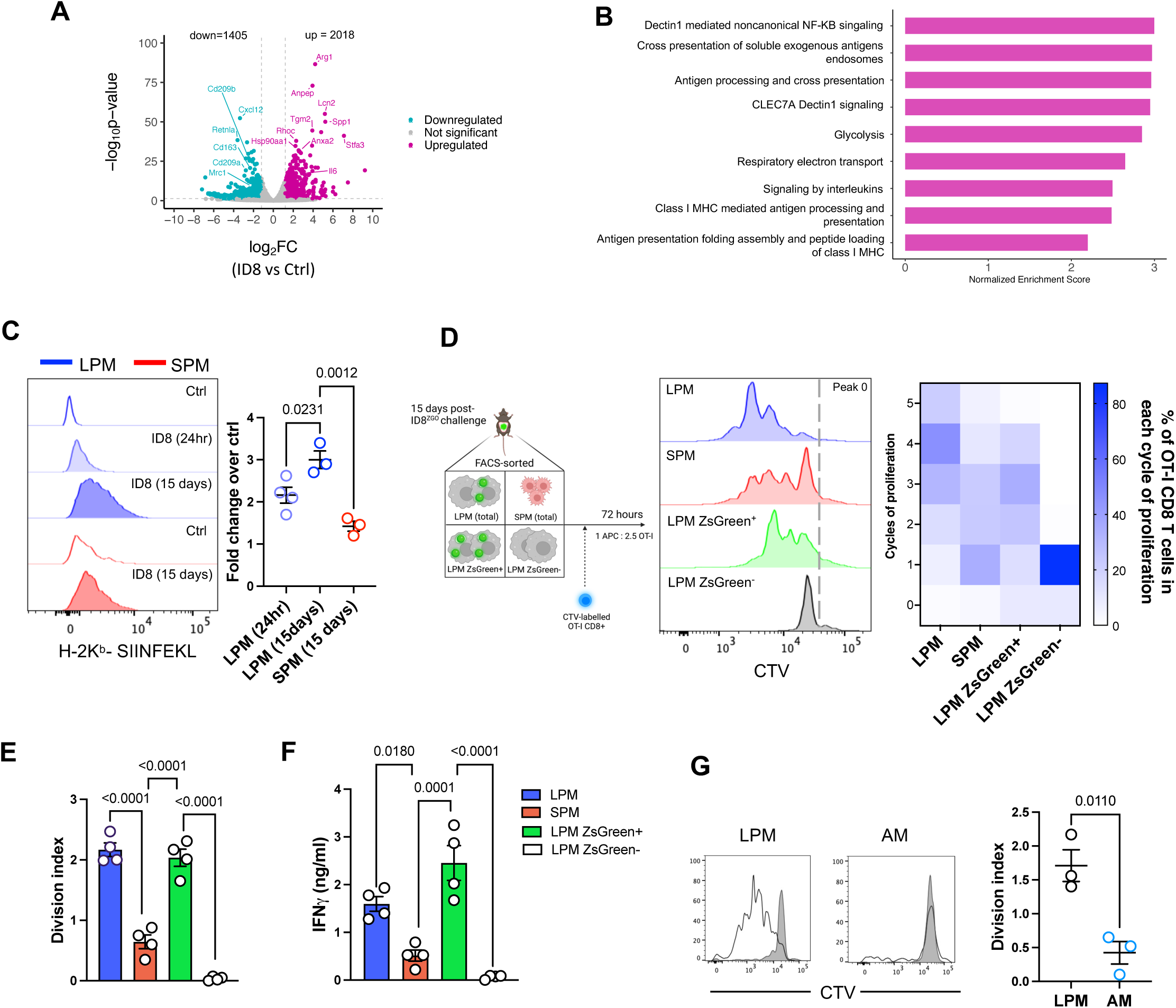
Uptake of cancer cells by LPM induces transcriptional remodeling and cross-presentation of tumour-derived antigens. **A)** Bulk RNA seq of cell-sorted tumor loaded (LPM-ID8) or control (LPM Ctrl) was performed 24 hr after challenge. Volcano plot showing genes upregulated (magenta) and downregulated (cyan) in LPM-ID8 vs control (log2Fold change >1.5 and <-1.5 respectively, p value > 0.05). **B)** Gene Set Enrichment Analysis (GSEA) was performed against Reactome gene sets of control LPM and LPM-ID8. NES: Normalised Enrichment Score. **C)** Histograms show labelling by 25D1.16 (specific for MHC-I-OVA complex) on LPM and SPM isolated 24 hours and 15 days after challenge with ID8-OVA and the corresponding quantification (fold change over control). Data represent mean ± SEM of one of two independent experiments (n= 3 to 4 mice). One-way ANOVA followed by Dunnett’s multiple comparisons test. **D)** LPM, SPM, LPM ZsGreen^+^ and ZsGreen^-^ were sorted 15 days after ID8^ZGO^ injection and co-cultured with CTV labeled naïve OVA specific CD8 T cells (OT-I), as depicted in the scheme. Primary histograms and heat map quantification of the fraction of proliferated OT-I in each cycle of division. **E)** OT-I proliferation plotted as division index. **F)** IFNγ secretion release by OT-I cells after 72 hours of co-culture. (D,F: n=4; each point is pooled from 2 animals). Data represent mean ± SEM, one-way ANOVA followed by Šídák’s multiple comparisons test. **G)** LPM and lung alveolar macrophages (AM) were loaded with apoptotic ID8^ZGO^ and co-cultured with CTV labelled OT-I. Histograms show proliferation profiles induced by control LPM or AM (filled) or ID8^ZGO^ pulsed cells (empty) and the corresponding division index. n=3, Data represent mean ± SEM. Statistical analysis was conducted by unpaired t-test.

### TIM4 deficiency impairs uptake and cross-presentation of tumor associated antigens by LPMs

To directly examine the role of TIM4 during cancer cells uptake and cargo processing, we next injected ID8^ZGO^ in WT and TIM4 null recipients (Tim4 KO) ^33^. The immune peritoneal compartment was analyzed 15 days after challenge. The influx in SPM and the frequency of LPM were equal between genotypes (**Fig3 A, B**). Proportions of cDCs, T cells, monocytes and neutrophils were comparable, except for a slight decrease in neutrophils and an increment in the fraction of CD4 T cells in knockout animals (**Fig S4A**). Cancer cell uptake by peritoneal phagocytes was similar between genotypes, with the exception of LPM. Indeed, the fraction of ZsGreen^+^ LPM was reduced from 67.75 % (± 9.50) in WT to 38.34%(± 7.14) in TIM4 KO, indicating that optimal engulfment of cancer cells by LPM requires TIM4 and alternative receptors cannot fully compensate (**Fig 3C, S4B**). TIM4-blocking antibodies resulted in a similar reduction of phagocytosis by LPM, confirming a receptor-specific effect (**Fig S4C**). To understand how entry via TIM4 may affect subsequent cellular reprogramming, we next compared the transcriptional profiles of steady-state and ZsGreen^+^ WT and TIM4 KO LPM. As a control, we verified that scavenger and phagocytic receptors were similarly expressed in the two genotypes, excluding compensatory effects (**Fig S4D**). Gene profiles at steady state were relatively comparable between genotypes, showing some changes in genes belonging to mitochondrial respiration (enriched in WT) (**Fig S4E,F, Supplementary table 2**). In contrast, the uptake of tumor cells induced a different response in the two genotypes (WT vs WT-ID8 and TIM4 KO vs TIM4KO-ID8, **S3G vs S2A**). Consistently, a direct comparison between WT and TIM4 KO LPM containing phagocytosed cancer cells showed differential expression of genes and biological processes (**FigS4H,3D**). Specifically, uptake induced enrichment of genes regulating cellular metabolism (cellular respiration, glycolysis, oxidative stress) in WT cells, whilst cholesterol biosynthesis and lipid metabolism were depleted with respect to TIM4 deficient cells. Notably, processes implicated in cytoskeletal dynamics, vesicular trafficking, inflammatory response and antigen processing/presentation were also enriched in WT LPMs (**Fig3D, Supplementary table 2**). Genes defining these processes include kinesins, Rho GTPases, actin nucleation-promoting factors, interferon and cytokine signalling, proteasomal components, membrane transport and MHC class-I molecules. Of interest, genes governing cholesterol efflux and lipid metabolism were preferentially enriched in TIM4 KO cells (**Fig 3E**). Collectively, these data suggest that engulfment via TIM4 induces a proinflammatory transcriptional reprogramming of LPMs that includes upregulation of the machinery for antigen presentation. When TIM4 is absent, residual uptake via alternative routes triggers lipid reprogramming and cholesterol efflux, which is consistent with a decreased inflammatory profile.

**Figure 3.**
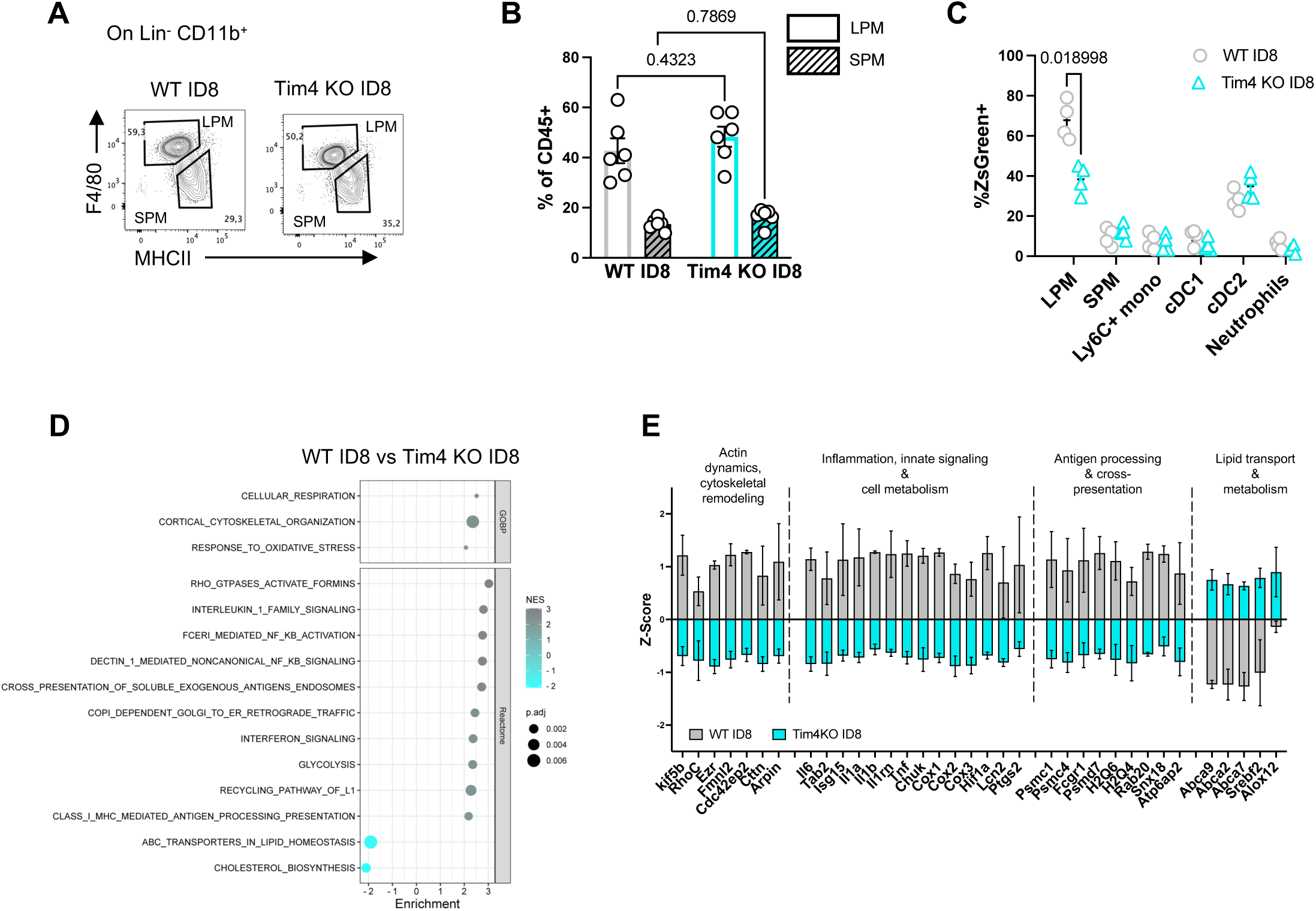
Tim4 controls transcriptional remodeling of LPMs. **A-B)** WT and Tim4 KO mice were injected intraperitoneally with ID8^ZGO^ and peritoneal cells were isolated after 15 days. **A)** Representative flow cytometry dot plots of SPM and LPM in the peritoneal cavities of WT and Tim4 KO. n=4, Data represents mean ± SEM from one of the three independent experiments. two-way ANOVA followed by Šídák’s multiple comparisons test. **C)** The phagocytic index is plotted as the fraction of ZsGreen^+^ cells within total cells in each of the indicated populations. Data are mean ± SEM of one of two independent experiments (n=4; multiple t-test followed by Welch correction. **D)** GSEA was performed against GO and Reactome gene sets of WT and Tim4 KO LPM-ID8. The dot plot shows selected pathways, significantly enriched in WT LPM-ID8. NES: Normalised Enrichment Score. **E)** Distribution of Z-scores of genes defining the enriched pathways in WT-ID8 related to four indicated processes.

### TIM4 associates with F-actin in nascent phagosomes in LPMs

To extend transcriptional data to cellular phenotypes, we established ex-vivo assays to examine phagocytosis and intracellular trafficking of cargo by confocal imaging and functional analysis, using apoptotic thymocytes as a surrogate phagocytic target. LPM isolated from resting WT or Tim4 KO mice were pulsed with CFSE-labelled apoptotic thymocytes for 15 min, washed and chased for a further 15 or 30 min before fixation and labelling. We ranked apoptotic-LPMs interactions across three stages: stage 0, corresponding to cell-cell proximity without any evident engagement; stage I, corresponding to conjugate formation and phagocytic cup formation; stage II, corresponding to phagosome closure (**Fig 4A**). After 15 min, WT cells were equally distributed between stage I and stage II and further progressed toward stage II after 30 min. As expected, a large fraction of Tim4 null LPMs were blocked in stage I at both time points, in line with impaired phagocytosis. Nevertheless approximately 27% and 32% had progressed to stage II, after 15 and 30 min respectively (**Fig 4B**). Careful inspection of the images showed a prominent cell membrane curvature enriched in TIM4 at the phagocytic cup, overlapping with recruited F-actin (**Fig 4 A, C, D**). In addition, recently formed phagosomes in stage II were surrounded by a sharp ring marked by TIM4, coincident with a circular F-actin ring. In contrast, Tim4 null LPMs accumulated a thick F-actin interface at the site of phagocytic interaction and cells already internalized were mostly devoid of surrounding actin (**Fig 4 A, E**). This last result aligns with findings in zebrafish microglia where the actin layer forming around internalized phagosomes depends on TIM4 ^34^. Thus, consistent with transcriptional data and previous studies in other cellular models ^35^, TIM4 drives actin remodeling at the phagocytic cup and travels with cargo during internalization in LPM.

**Figure 4.**
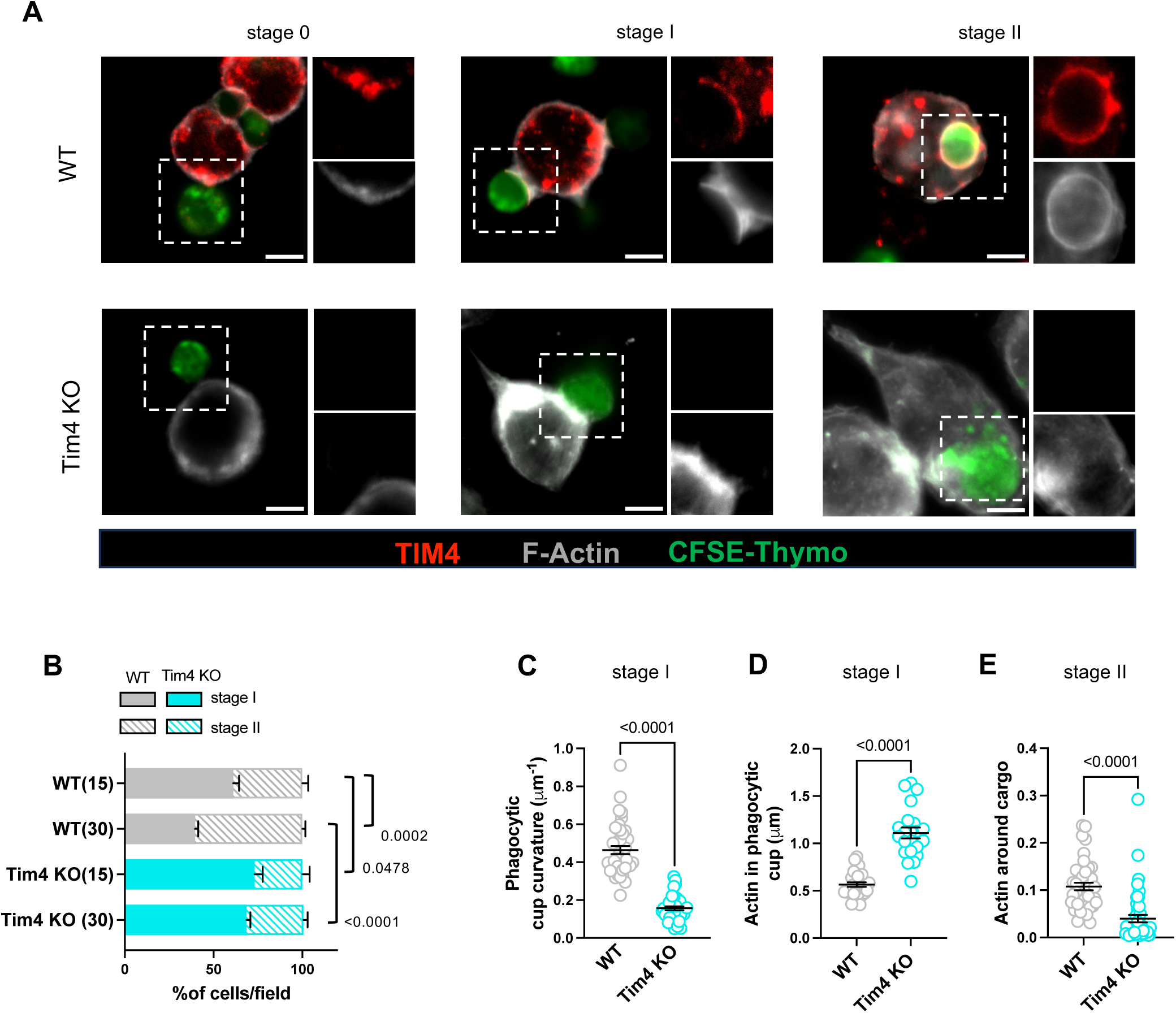
Tim4 localizes with actin at the phagocytic cup and it is internalized in nascent phagosomes. **A)** WT and Tim4 KO LPM isolated from naïve mice were pulsed for 15 minutes with CFSE-labelled apoptotic thymocytes followed by chase up to 30 minutes. After fixing, cells were stained using antibodies to TIM4 and phalloidin to label F-Actin. Representative confocal images show the three stages of phagocytosis used for classifying the interactions. Scale bar=5μm. **B)** Quantification of the number of events in stage I and stage II in WT and Tim4 KO LPM at the the indicated time points. Data are representative of n>30 events/fields containing >15 LPM each, pooled from three independent experimental replicates. Data represent mean ± SEM, two-way ANOVA followed by Šídák’s multiple comparisons test. **C)** Curvature index of the phagocytic cup calculated on cells in stage I. **D)** Thickness of F-Actin in phagocytic cup in μm. n>15 cells, data represent mean ± SEM. Statistical analysis was conducted by unpaired t-test. **E)** F-Actin integrated density in nascent phagosome around the cargo normalised with F-Actin density within each cell. n>20 cells, data represents mean ± SEM. Statistical analysis was conducted by unpaired t-test.

### TIM4 controls the kinetics of degradation of ingested cargo

To examine subsequent steps of cargo progression, we selected cells in stage II to normalize for defective uptake. At 15 min, intracellular cargo was included in vesicles negative for lysosomal markers which started to accumulate LAMP-1 signal after 30 min. After 45 min Internalized apoptotic cells were mostly fragmented and the remaining intracellular green signal was entirely contained within the lumen of lysosomal vesicles (**Fig 5 A,C**). Surprisingly, Tim4 KO LPM, accumulated a dense crowd of LAMP-1^+^ vesicles around ingested cargo already after 15 min of chase, showing signs of incipient cargo fragmentation (**Fig 5B,C**). After 30 min, majority of engulfed apoptotic cells were degraded into smaller particles and green signals had started to fade by 45 min. Consistently, scoring the size of cargo as intact, partially or fully fragmented, revealed a significantly faster accumulation of degraded material within Tim4 KO cells (**Fig5 D**). The same trend was observed when feeding WT or Tim4 KO LPMs with apoptotic ID8, suggesting that TIM4 controls downstream trafficking independently of the cargo nature (**Fig S5 A,B**). To reinforce this result, we set up two complementary approaches to quantitatively assess the rate of cargo degradation. In the first, dying thymocytes were loaded with cypher, a pH sensitive probe that emits fluorescence upon acidification in acidic organelles (**Fig S5C**) and CTV, as a control for uptake. WT or Tim4 KO LPMs were fed with labeled apoptotic thymocytes ex-vivo and cells were analyzed by flow cytometry to follow the kinetic of acidification. As shown in **Fig 5E**, fluorescence intensity steadily increased over time in both genotypes. However, in Tim4 KO cells, the first burst in fluorescence emission occurred after 5 min and acidification was consistently higher than in WT cells, at each time point analyzed. Second, to directly assess cargo degradation we prepared liposomes (lipo) containing phosphatidylserine (PS) to target TIM4, or phosphatidylcholine (PC) as control (**Fig S5D**). DQ-BSA (a self-quenched conjugate that emits bright fluorescence only upon proteolytic cleavage in endosomes) and BSA-Alexa 647 as control, were encapsulated into PS-and PC-lipo. We verified that PS-Lipo were specifically taken up by TIM4^+^ LPM (**Fig S5E**). TIM4^+^ LPMs having internalized lipo-PS started to degrade DQ-BSA 15-20 min after internalization and reached a plateau after 75 min.

**Figure 5.**
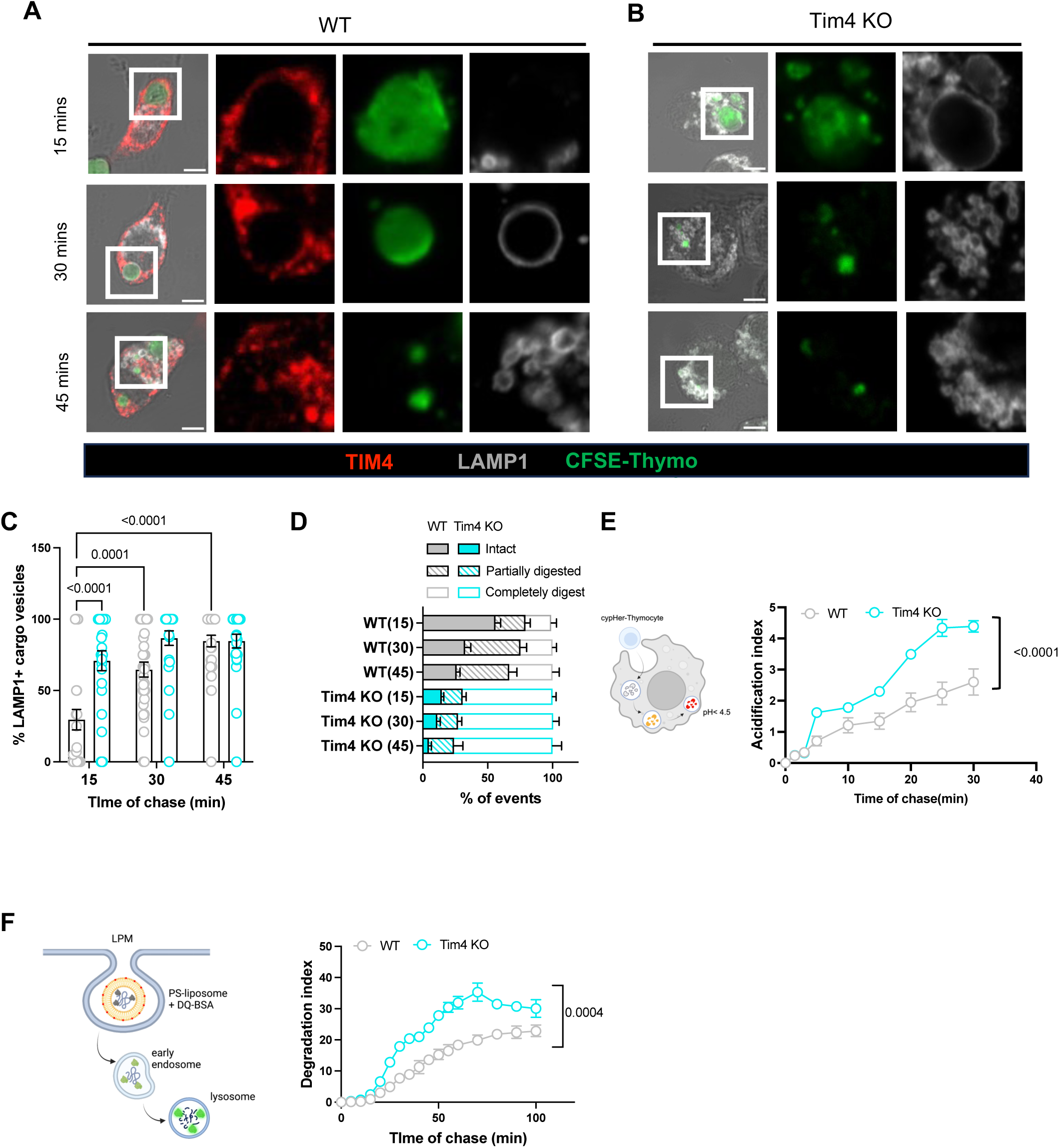
Tim4 controls phagosome maturation and cargo degradation. **A,B)** Representative confocal images showing lysosomes (LAMP-1) and TIM4 labelling in WT and Tim4 KO LPM pulsed with CFSE-labelled thymocytes and chased for 15, 30 or 45 min. Scale bar=5μm. **C)** Bars show the fraction of internalized cargo associated to LAMP1 vesicles. A minimum of 20 cells in 3 independent experiments, one of three independent experiments is represented (mean ± SEM). Two-way ANOVA followed by Šídák’s multiple comparisons test. **D)** The fraction of intact, partially digested or completely digested cargo withing cells that have internalized was quantified for each of the indicated time points. Data represent >10 fields consisting at least 15 cells per time point, per group. **E)** WT or Tim4 KO LPM were pulsed for 15 min with apoptotic thymocytes labelled with cypHer and CTV. The fluorescent intensity of cypHer was measured on CTV^+^ F4/80^+^ LPM every 5 minutes, during the following 30 min. The acidification index is calculated as fold change of cypHer fluorescent intensity relative to time 0 min. n=3 independent experimental replicates, data represent mean± SEM. Statistical analysis was conducted by comparing slopes of two groups by simple linear regression. **F)** LPM were pulsed for 15 minutes with 2μM liposomes (PC-DQ BSA or PS-DQ BSA)/100,000 cells for 15 minutes. Cells were washed with PBS to remove any free liposomes. Median fluorescence intensity of DQ BSA was acquired every five minutes up to 100 minutes. Fold change of DQ BSA median fluorescent intensity over time 0 min was plotted as degradation index. n=3 independent experiments, data represents mean± SEM. Statistical analysis was conducted by comparing slopes of two groups by simple linear regression.

Remarkably, fluorescent emission by DQ-BSA started earlier in TIM4 KO cells and was significantly higher at every time point (**Fig5F**), indicating faster and larger degradation in TIM4 KO cells. In summary, we conclude that entry via TIM4 in LPM orchestrates the progression of ingested cargo and balances phagosome maturation, preventing immediate antigen degradation.

### Tim4 is required for cross-presentation of cancer associated antigens by LPMs

Finally, we performed functional assays to link the above evidence to cross-presentation and induction of anti-tumoral responses in the context of WT and Tim4 KO LPM. Labeling LPM with pMHC-I OVA specific antibodies 15 days after challenge showed a consistent reduction in TIM4 KO (**Fig 6A**). Next, we plated total or ZsGreen^+^ LPM with OT-I to assess proliferation and IFN-γ release. In line with previous results (Fig 2D), WT cell induced several cycles of proliferation and consistent IFN-γ release. In contrast, total or ZsGreen^+^ TIM4 null LPM triggered poor CD8 T cells proliferation and IFN-γ production, indicating blunted cross-presentation and reduced induction of effector T cell functions (**Fig 6B,C**). Interestingly, immune profiling of the peritoneal infiltrate 15 days after challenge revealed an accumulation of functional, tumor-specific, endogenous CD8 T cells in WT mice, which was significantly reduced in Tim4 KO animals (**Fig 6D, E**). Furthermore, we confirmed the superior ability of WT LPM to cross-present tumor antigens immediately after cancer cells encounter (24 hr, **Fig 6F-H**), ruling out differences that may accumulate in TIM4 KO animals during initial tumor growth. Importantly, tumor specific CD8 T cells transferred in the peritoneum of animals that had been challenged with ID8^ZGO^ 24hr before, started to proliferate and produce IFN-γ only in the context of WT recipient and not in TIM4 KO host (**Fig 6 I**). Finally, we explored the potential of artificial membranes exposing the TIM4 ligand PS to deliver tumors antigens to cross-presenting macrophages. To this end, whole OVA protein was encapsulated within PS-liposomes or, as control, PC-liposomes. Mice were intraperitoneally injected with PS or PC-OVA liposomes and co-transferred with CTV-labelled OVA specific CD8 T cells to track activation. As shown in **Fig 6J**, PS-OVA induced robust proliferation of OT-I cells, while PC-OVA liposomes or PS-liposomes containing a control protein induced little or no T cell proliferation.

**Figure 6.**
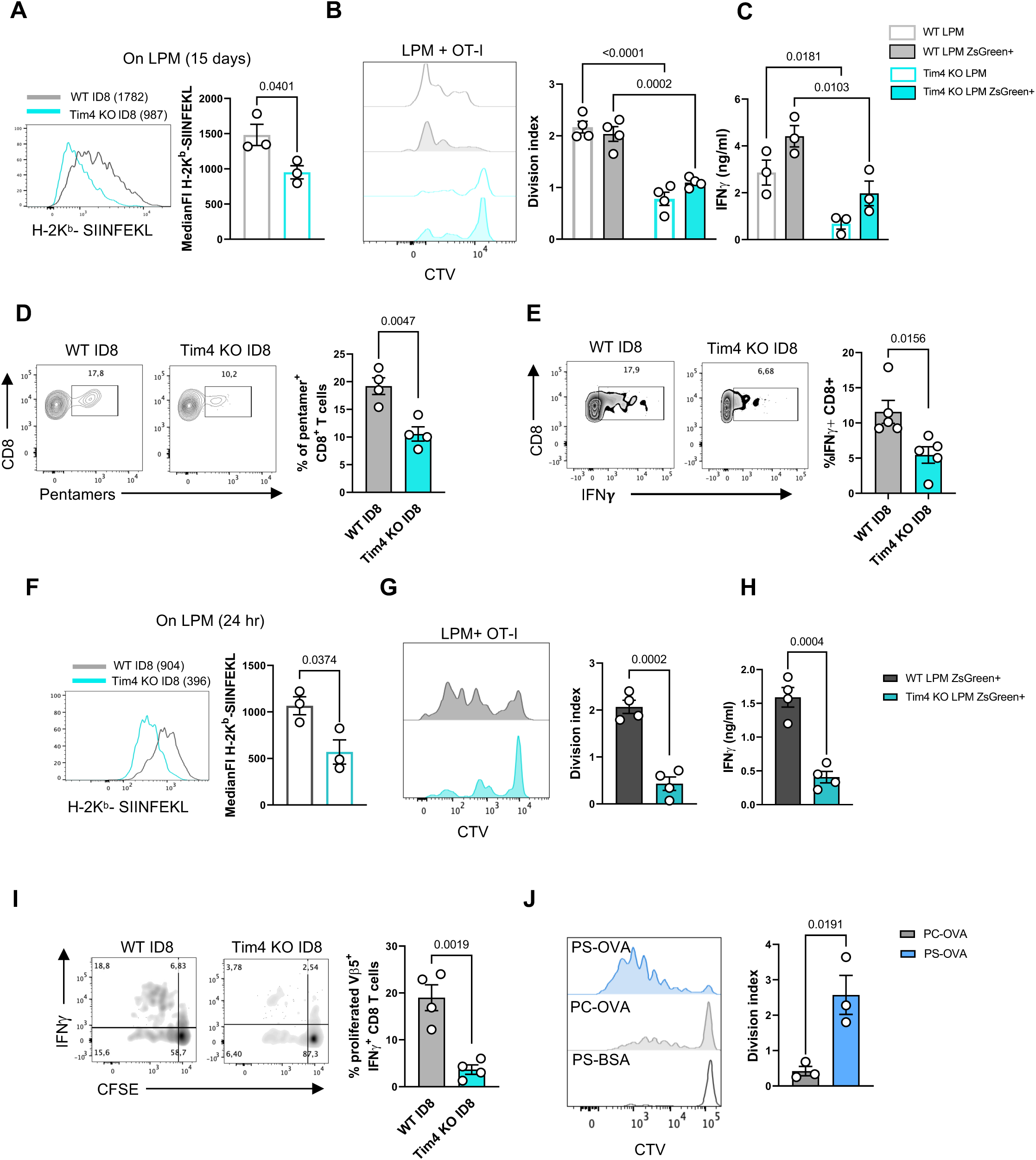
Anti-tumoral CD8 T cell responses to peritoneal metastasis depends on cross-presentation by Tim4 LPMs. **A)** Histograms show MFI of WT and TIM4 KO LPM stained with 25D1.1.6 antibody, 15 days after ID8^ZGO^ tumor induction. n=3, data represent mean±SEM from one of the two independent experiments. Statistical analysis was conducted by unpaired t-test. **B)** Representative histograms and index of OT-I proliferation induced ex-vivo by total or ZsGreen^+^ LPM sorted from WT and Tim4 KO, 15 days after challenge. **C)** IFNγ production in the supernatant of LPM-OT-I CD8 T cell co-cultures as measured by ELISA. n=4, data represent mean±SEM, two-way ANOVA followed by Šídák’s multiple comparisons test. **D-E)** In vivo activation of peritoneal tumor-specific CD8 T cells (identified by OVA-specific Pro-5 pentamers) in WT and Tim4 KO animals, 15 days after challenge. Representative dot plots and quantification, n=4 animals per group. Data represent mean±SEM, unpaired t-test. **E)** Intracellular production of IFN-g by endogenous CD8 T cells restimulated ex-vivo with OVA class-I peptide. Representative dot plots and quantification. n=5, data represent mean mean± SEM. Statistical analysis was conducted by unpaired t-test. **F)** 25D1.1.6 labelling of LPM 24 hr after tumor challenge (data are mean±SEM from one of two independent experiments, n=3 mice each, t-test). **G-H)** As in B), cells were harvested 24 hr after ID8^ZGO^ challenge. **I)** WT and Tim4 KO mice were injected with CFSE-labelled OT-I CD8 T cells and challenged, 18 hours later, with apoptotic ID8 OVA, intraperitoneally. Peritoneal cells were harvested after 48 hours to assess OT-I CD8 T cell proliferation and IFNγ production by intracellular labelling. n=4, data represent mean±SEM from one of the two independent experiments, unpaired t-test. **J)** WT mice were injected with CTV-labelled OT-I, followed by intraperitoneal injection of PC-OVA, PS-OVA and PS-BSA liposomes, 18 hours later. CD8 T cell proliferation and quantification of proliferation index. n=3, data represent mean± SEM from one of the two independent experiments. Statistical analysis was conducted by unpaired t-test.

In conclusion, these results demonstrate that targeting of TIM4 on LPM, by cancer cells or artificial membranes, promotes immediate engagement of anti-tumoral CD8 T cells against incoming metastasis in the peritoneum.

## Discussion

Understanding the specific function of different classes of tumor-associated macrophages at different stages along tumor progression is key for developing novel macrophage-centered therapeutic axis. Here we demonstrate that, shortly after seeding in the peritoneum, LPMs can transiently cross-present tumor antigens. This exquisite function of LPM is supported by the PS receptor TIM4 that maximizes internalization of cancer cells, regulates transcriptional circuits induced by engulfment and orchestrates phagosomal maturation to promote cross-presentation of tumor antigens.

The role of LPM in cancer has been addressed mostly in the context of antigen presentation-independent functions and at late stages of tumor progression. Consistent reports have shown that tissue-resident LPMs of embryonic origin proliferate in situ and progressively incorporate inflammatory monocytes under tumor-derived cues ^22,27,32,36^. In parallel, LPM shift from a tumoricidal, high-inflammatory profile at early stages to an anti-inflammatory profile in advanced tumors. The latter is characterized by high oxidative phosphorylation and glycolysis, enhanced mitophagy, altered lipid metabolism, increased cholesterol efflux and dominance of IL-4 signaling over IFNγ signaling ^20–22^. Our data, focusing on very early events of metastatic colonization of the peritoneum, unveil a distinct side of LPM. Indeed, cancer cell recognition by naïve LPMs triggers vigorous uptake, followed by an inflammatory and metabolic response that culminates in the presentation of ingested antigens on MHC class-I. The induction of inflammatory genes program also indicates that the uptake of cancer cells, unlike normal apoptotic cells ^37^, is not silent and may actually contribute to remodeling of the compartment, as it has been recently described in monocyte-derived macrophages expressing TREM2 ^38^. Interestingly, the capacity of LPM to cross-present was strongly inhibited in late tumors, which is consistent with an active educational mechanism that shapes resident macrophages towards a pro-tumoral phenotype.

Cross-presentation of cancer antigens was previously described for CD206^+^ monocytes derived cells ^13^, upon CD47 blockade in bone marrow-derived macrophages ^39^, in red pulp splenic macrophages ^40^ and in lymph node resident CD169^+^ cells ^14,41^. So far, intrinsic cross-presentation by a well-defined population of embryo-derived cells such as LPM is unprecedented and suggests a new layer of cancer immune surveillance, at sites of metastatic colonization.

Several cellular adaptations have been characterized, especially in dendritic cells, that enable cross-presentation by phagocytes. First, receptors that mediate the uptake of dead cells such as Clec9-DNGR-1, CD36, Axl, SCARF-1 and TIM3 have been suggested to drive cross-presentation of ingested antigens in dendritic cells ^42–47^. Second, depending on the cellular context, cross-presentation is facilitated by decreased acidification in phagosomes to preserve antigen integrity; active transfer to the cytosol to intersect the conventional MHC class-I pathway and specialized pathways of vacuolar trafficking ^42,48–51^.

In macrophages, cross-presentation is proposed to occur mainly via the vacuolar pathway, i.e., loading of peptides would occur within the lumen of phagolysosomes ^52,53^. Based on the finding of accelerated antigen degradation in knockout cells, we speculate that TIM4, in cooperation with accessory signalling receptors ^54^, may be essential for the biogenesis of and/or routing to a compartment specialized for cross-presentation. Remarkably, a recent study showed that grafting TIM4 on engineered T cells confers them with the ability to cross-present ^55^.

Conflicting with our findings, previous reports suggested that TIM4 expression by TAM and bone marrow-derived macrophages controls LC3-mediated phagocytosis, a form of non-canonical autophagy that accelerates antigen degradation, blunting cross-presentation ^28,56^. This discrepancy may depend on the different cell types analyzed (mono and BM-derived macrophages vs resident LPM), suggesting a context-dependent function of the receptor. It is also important to note that we and others (Roberts et al., 2023, Dransfield et al., 2015), could not detect TIM4 expression in bone marrow-derived macrophages.

Interestingly, our transcriptomic analysis revealed a relatively higher expression of several genes involved in lipid transport in TIM4 null macrophages (**Fig 3E, Fig S4H**). This includes members of ABC family such as *Abca2* and *Abca7*, known as lipid transporters^57^, Abca9, a speculated mediator of lipid homeostasis^58^ as well as Apoe, which reportedly drives lipid transport and immunosuppression via induction of *Cxcl1* and *Cxcl5*^59^. This transcriptional profile hints towards increased lipid export, which would be consistent with the observed decreased induction of inflammatory genes^32,60^. A further intriguing link between TIM4 and lipid metabolism recently emerged in adipose tissue macrophages, where the receptor was shown to mediate the transfer of scavenged lipids into lysosomes, promoting lysosomal functions and ABCA1-mediated post-prandial cholesterol transport^26^. Thus, establishing the precise causal link between TIM4 regulation of lipid metabolism and cross-presentation remains an interesting area of future investigations.

The ability of TRM to cross-prime CD8^+^ T cells and drive differentiation into polyfunctional effectors has been documented before in the context of breast cancer. Importantly, macrophages mediating this function display high TIM4 expression and are associated with better prognosis in patients ^8^. In the present study, we also observed immediate activation of tumor-specific CD8^+^ T cells by TIM4 macrophages and accumulation of cytotoxic CD8^+^ T cells at day 15, which were significantly impaired in TIM4 null animals. Our observation, however, remains strictly restricted to early stages of tumorigenesis. At late stages, the peritoneal cavity is heavily infiltrated by mono-derived macrophages, which instead mediate exhaustion and deactivation of T cells ^27^. Therefore, cell-intrinsic changes in LPMs, combined to massive infiltration of immune suppressive mono-derived cells, collectively override immunosurveillance. Currently, the unavailability of tools to conditionally ablate Tim4^+^ on LPM at defined time points precludes evaluation of later outcomes.

In summary, our findings reveal an important intracellular function of TIM4 that, besides mediating uptake of PS-expressing cells, contributes to channel antigens for cross-presentation of ingested antigens, enabling cancer immune surveillance at initial stages.

## Acknowledgements

This work was supported by AIRC IG 21635 to FB. SJ, LLR and LGM were supported by ICGEB pre and post-doctoral fellowships. RA is supported by Italian Telethon. We thank Giulia Canarutto and Silvano Piazza (ICGEB Trieste) for the initial RNA-seq data processing. We also extend our gratitude Dr M.Ruhland and Prof. M. Krummel for providing reporter plasmids for ZsGreen and OVA and Dr Maria Casanova-Acebes (CNIO, Madrid) for insightful discussions on the project.

## Author contributions

FB and SJ conceived the study. FB supervised the planning and execution of experiments. SJ performed experiments and data analysis. LLR, LGM, GMP and RA assisted in the execution of experiments, figure preparation and critical discussion of the data. LGM also curated biostatistics. MP and MG prepared lipid particles for PS targeting.

## STAR Methods

### RESOURCE AVAILABILITY

- Lead contact Further information and requests for resources and reagents should be directed to and will be fulfilled by the lead contact, Federica Benvenuti. Institutional and funding agency requirements for resource and reagent sharing will be followed.
- Materials availability This study utilized derivatives of ID8 cell lines namely ID8-ZsGreen, ID8^ZGO^ and ID8- OVA that were generated in our lab. This resource is available upon request to the lead contact as indicated above.
- Data and code availability
  - Sequencing data have been deposited at GEOdatabase (GEO:GSE242989), which are publicly available as of the 16/09/2023.
  - This paper does not report original code.
  - Any additional information required to reanalyze the data reported in this paper is available from the lead contact upon formal request.

### EXPERIMENTAL MODEL AND SUBJECT DETAILS

- Mouse models C57BL/6, OT-I (C57BL/6-Tg(TcraTcrb)1100Mjb/J) and Balb/C mice were purchased from Envigo or Jackson Laboratories, respectively. Timd4^−/−^ (Tim4 KO) mice ^33^ were a kind gift of Professor S. Nagata (Osaka University). Animals were maintained in sterile isolators at the ICGEB animal Bio-experimentation facility (12 h/12 h light and dark cycle, 21°C ± 2°C). Animal care and treatment were conducted with national and international laws and policies (European Economic Community Council Directive 86/609; OJL 358; December 12, 1987). All experiments were performed in accordance with the Federation of European Laboratory Animal Science Association (FELASA) guidelines. The study was approved by International Centre for Genetic Engineering and Biotechnology (ICGEB) board for animal welfare and authorized by the Italian Ministry of Health (approval number 459/2022-PR).
- Cell lines and cell culture The ID8 cell line was kindly provided by Elena Lacchetti, c/o Mario Colombo lab, Fondazione IRCCS Istituto Nazionale dei Tumori di Milano. To generate ID8 variants with fluorescent reporters and model antigen, vectors pSiren-ZsGreen and pCAGGS-ZsGreen-minOva ^31^, were provided by Dr Megan Ruhland. Briefly, ID8 cells were genetically engineered to stably express ZsGreen using viral transduction with ZsGreen construct. To make ID8^ZGO^ cells, ID8 parental cells were transfected with ZsGreen-minOVA construct and sorted using ARIA II sorter after every 2 passages until stable integrants were obtained. To generate ID8 OVA, ID8 cells were transduced with lentiviral vector pDual-liOva-puroR encoding liOva (kindly provided by David Escors Murugarren, CIB, Navarra). After puromycin selection, cells were subcloned and single cell clones were tested for HA expression by intracellular staining using anti-HA antibody (clone 3F10, Roche). The parental ID8 cells and the derivatives were maintained in DMEM media supplemented with 4.5 g/mL glucose (Invitrogen) and 10% fetal bovine serum (FBS, Euroclone) and Gentamicin (50 µg/mL, Gibco) at 37°C in 5% CO_2_ and were routinely tested for mycoplasma contamination. Cells were expanded to passage 3 and stored in aliquots in liquid nitrogen. Cells used for *in vivo* challenge have been passed less than five passages.

### METHOD DETAILS

- Establishment of peritoneal tumors: To track early events of peritoneal tumor establishment at day 15, exponentially growing ID8^ZGO^ or ID8 OVA cells were treated with trypsin (Gibco) and prepared as a single-cell suspension at a concentration of 2⍰×⍰10^7^ cells/mL in PBS. WT and Tim4 KO littermates from TIM4 colony were injected intraperitoneally with 100⍰μL of the cell suspension (2⍰×⍰10^6^ cells). To track post-phagocytic events at 24 hours, ID8^ZGO^ or ID8 OVA were exposed to UV-C in a UV cross-linker for 10 minutes, followed by 3 hr incubation at 37°C. Cells were then harvested by scraping and single cell suspension was prepared of density 3×10^6^ cells/1mL PBS. 100μL of this suspension (3×10^5^ cells) were injected in WT or Tim4 KO littermates. For all experiments, 100⍰μL of PBS was injected as a negative control in a parallel group of control mice. In TIM4 blocking experiments, mice inoculated with ID8^ZGO^ cells were treated with 125 μg (diluted in PBS) of anti-Tim4 (clone RMT4-53, InVivoMab) or isotype control (rat IgG2b,κ, clone LTF2,InVivoMab) at day -1, and later every 72 hours until the experimental endpoint at 15 days. At the experimental endpoint (24⍰h, 14⍰or 60 dpi) of each assay, mice were euthanized by cervical dislocation.
- Isolation of murine peritoneal and omental cells: To isolate peritoneal cells from control or tumor bearing mice, 6 mL of ice cold PBS was injected with syringe in the peritoneal cavity immediately after sacrifice, peritoneal wall was massaged for 2 minutes and cells were collected by syringe, followed by one wash with PBS and then proceeded for flow cytometry analyses. Omenta excised from control or ID8^ZGO^ injected animals were mechanically cut into small pieces and digested with Collagenase type 2 (265⍰U/mL; Worthington) and DNase (250⍰U/mL; Thermo Scientific) at 37⍰°C with gentle shaking for 30’. Collagenase action was then stopped by EDTA 10⍰mM (Invitrogen^TM^, Life technologies), and the cell suspension was filtered using 70⍰μm cell strainer (Corning). The single cell suspension thus obtained was then processed for flow cytometry analyses.
- Flow cytometry: The antibodies used for the experiments are listed in Key resources table and panels used to identify subsets are illustrated in Supplementary Table (**S1A, S2A, S3C**). Viability was assessed by staining with LIVE/DEAD^TM^ Fixable Aqua Dead Cell Stain Kit (Invitrogen^TM^) Life Technologies). For cell staining, FcR binding sites were blocked by using αCD16/CD32 (Clone 93, Biolegend). Samples were then stained with specific antibodies in⍰1% BSA in PBS and fixed with ⍰1% PFA in PBS. Production of IFNγ in CD8 T cells isolated from tumor-bearing mice after restimulation was analysed by intracellular staining of IFNγ in cells fixed and permeabilized using Fixation/Permeabilization solution kit (BD Biosciences) following manufacturer’s instructions followed by IFNγ staining. For intracellular staining of OVA-HA, ID8 OVA cells or lung cell suspensions were fixed and permeabilized using Fixation/Permeabilization solution kit (BD Biosciences) following manufacturer’s instructions, and then stained with rat monoclonal anti-HA (3F10, Roche) and then with anti-rat A647 (Thermo Fisher). Absolute cell count was performed by adding CountBright^TM^ Absolute Counting beads (Invitrogen) to the samples following manufacturer’s instructions. After dead cell exclusion and exclusion of cell doublets, the subsequent populations were identified according to the gating strategy provided in Supplementary Fig S1A, S2A, S3C. Phosphatidylserine present on ID8 and derivative cells after UV exposure or after isolation from tumor injected mice at day 15 or in thymocytes after dexamethasone treatment was assessed by Annexin V/7AAD kit (Biolegend). For sorting specific macrophage subsets, suspension of labelled macrophages was prepared in 2% FBS in PBS, and cells were sorted using FACSAria II. Flow data were acquired with FACS Celesta (BD Biosciences) and analyzed with Diva software (BD Bioscience) or FlowJo software (Tree Star, Inc.).
- Induction of apoptosis: To induce apoptosis in ID8 and derivative cells, cells were exposed to UV-C with UV lamp (254 nm, 6W, output 1.7W) for 10 minutes followed by 4 hours incubation in PBS at 37°C. Cells were scraped to detach and prepare a single cell suspension and were stained for AnnexinV/7AAD as described above to analyze induction of apoptosis. Thymocytes were harvested from Balb/C animals and were incubated with dexamethasone (10 μM) in DMEM + 10% FBS medium for 4 hours, followed by three washes with DMEM + 10% FBS.
- Ex vivo priming of OT-I cell by cell sorted macrophages For ex-vivo T cell activation assay, LPM, SPM, LPM ZsGreen^+^ or LPM ZsGreen^-^ sorted by flow cytometry from ID8-tumor bearing mice and were plated in U-bottom 96 wells and co-incubated with CTV-labelled OT-I T cells at the ratio 1:2.5 (1×10^4^ macrophages and 2.5 × 10^4^ CD8^+^OTI). After 70⍰h of co-culture OTI T cell proliferation was assessed by flow cytometry. As positive control, 1×10^4^ sorted LPM were pulsed with class I OVA peptide (SIINFEKL, 90nM) for 2 hours. After 2 h stimulation, cells were washed twice in PBS and co-incubated with 2.5×10^4^ OT-I CD8 T cells. Division index was calculated using proliferation platform of FlowJo which is equal to the Total Number of Divisions / The number of cells at start of culture. IFNγ production was detected in the supernatant after by ELISA Max Standard sets (Biolegend), following manufacturer’s instructions.
- Ex vivo phagocytosis and cross-presentation by peritoneal and alveolar macrophages Peritoneal and alveolar macrophages were sorted from 8-weeks old WT mice according to gating strategy in (**Fig S1A, S3C**). 1.5×10^5^ cells were plated in 96-well U-bottom plates in IMDM + 10%FBS medium for 2 hours. Apoptotic ID8^ZGO^ were then added to at the ratio 1:5 (1.5×10^4^ macrophages and 7.5×10^4^ tumor cells). After 1 h of incubation, cells were washed thrice with PBS to remove non-internalized/non-attached tumor cells and either processed to asses phagocytosis by flow cytometry or co-cultured with CTV-labelled OT-I CD8+ T cells in the ratio 1: 2.5 (1.5×10^4^ macrophages to 2.75×10^4^ OT-I T cells). 72 hours later, proliferation of OT-I T cells was analyzed by flow cytometry. Division index was calculated using proliferation platform of FlowJo.
- Bulk RNA sequencing of ZsGreen+ LPM and analyses: LPM (F4/80^high^ MHCII^low^) were sorted from naïve WT and TIM4 KO mice, whereas, LPM ZsGreen^+^ (F4/80^high^ MHCII^low^ ZsGreen+) were sorted from WT ID8 and TIM4 KO ID8 mice. RNA from sorted cells was extracted using RNeasy micro kit (Qiagen) following manufacturer’s instructions. RNA purity, integrity and concentration were determined by NanoDrop (ND-1000 Spectrophotometer, ThermoFisher Scientific) and TapeStation 2200 (Agilent Technologies Inc., Santa Clara, CA, USA). Afterward, 100 ng of total RNA was used to prepare RNA libraries following the instructions of the SMART-Seq®mRNA LP, Takara bio kit. Library quality was assessed using TapeStation HS D5000 Screen tape (Agilent) and a narrow distribution with a peak size of approximately 400 bp was observed in all cases. Libraries were quantified by LightCycle qPCR and sequenced in a HiSeqX analyzer (Macrogen) in a run of 2 × 75 cycles and a high output sequencing mode. Approximately twenty million reads were obtained and analyzed for each sample. Fastq files coming from sequencing platform were merged and basic quality controls were performed with FASTQC and PRINSEQ tools. FASTQ files were aligned to the mm10 reference genome, reads were dereplicated for PCR duplicates and gene counts were generated using STAR v.2.5 using quantMode GeneCounts. Normalization of raw counts followed by differential gene expression analysis was performed with DESeq2 ^61^ package for R (version 4.1.2). Locally developed scripts were used to format and annotate the differential expression data output from DESeq2. Significant differentially expressed genes were identified as p<0.05, and log_2_foldchange <1.5 or >1.5.Gene Set Enrichment Analysis (GSEA, Broad Institute) was used to examine enriched pathways. The output of GSEA included an enrichment score (ES), a normalised enrichment score (NES) which accounts for the size of the genes being tested, a p value and an estimated false discovery rate or adjusted p value (p.adj). The input for GSEA was prepared by performing pairwise comparisons between DEGs from different populations. Comparative GO analysis was performed with clusterProfiler ^62^. Data is in the process of deposited in the NCBI’s Gene Expression Omnibus database (GSE242989).
- Preparation and characterization of PS and PC liposomes: The liposomes were prepared as described ^63^ from a lipid mixture (Avanti Polar Lipids) of phosphatidylserine (PS), phosphatidylcholine (PC), and cholesterol (CH) at 1∶1∶1.33 molar ratios, respectively (PS-presenting liposomes, PS-lipo). PS-lacking liposomes (PC-lipo) were prepared using from PC and CH (1.5∶1 molar ratio). The degree of PS exposure on liposomes was assessed by binding of FITC-AnnexinV to PS on liposome surface and analysis by FACS (**Fig. S5D**). Liposome preparations were characterized for the size using Zeta Plus particle size analyzer (Malvern Panalytical) and liposomes of size 130±15 nm were obtained. To enable detection of liposome uptake by macrophages and to assess rate of antigen antigen degradation after PS-mediated uptake, the liposomes containing Alexa Flour 647- Bovine Serum Albumin (BSA) and DQ-Bovine Serum Albumin were prepared. For targeted antigen delivery to LPM and tracking their cross-presentation ability liposomes containing protein OVA were prepared. Lipids (16 μmol) were transformed into the dried lipid film using rota-evaporator. During hydration of the lipid film, 500 μL equimolar solution of A647-BSA and DQ-BSA solution (1 mg⍰mL) or FITC-OVA (1 mg/mL) were added. To remove unencapsulated protein(s), liposomes were washed with saline, subjected to dialysis overnight and centrifuged (14000 × g, 40 min, 4°C). The final pellets were resuspended in PBS, to yield a final suspension concentration of 33.3 μmol lipid⍰mL. Encapsulation efficiency was determined by performing protein estimation assay (BCA) on the liposome solution.
- Degradation assay (DQ-BSA): WT and TIM4 KO LPM were pulsed for 10 minutes at 37°C with PS-liposomes or PC-liposomes encapsulating Alexa Flour 647–BSA and DQ BSA (2 μmol/million cells). Cells were washed twice with cold PBS in order to remove nonspecific binding /non-internalised liposomes from the cell surface. Cells were then acquired on FACS Celesta,BD for 90 min at every 3 min interval at 37°C and median fluorescent intensity of DQ-BSA normalized on Alexa Fluor 647-BSA^+^ cells was recorded DQ-BSA degradation index was calculated as log fold change of median fluorescent intensity of DQ-BSA on Alexa Flour 647^+^ F4/80^high^ macrophages over median fluorescent intensity of DQ-BSA on F4/80^high^ macrophages at time 0 min.
- Tracking the rate of lysosomal acidification using cypHer labelled thymocytes: LPM were isolated from peritoneal lavage with ice cold PBS from naïve, WT mice and allowed to adhere in μ-slide chambered coverslips (ibidi) for 6 hours. Thymocytes were isolated from Balb/C mouse and were incubated with dexamethasone (10 μM) in DMEM + 10% FBS for 4 hours at 37°C. After four hours, cells were washed twice with PBS and stained with CTV 5μM (Invitrogen) and cypHer5E-Alexa Fluor 647 5μM (Cytiva) in PBS + 10% carbonate buffer (pH 9.2) at 37°C for 20’. Unbound dye was quenched by adding complete medium and washing the cells. LPM were pulsed with apoptotic, labelled thymocytes for 10’ at 37°C and immediately stored on ice, followed by staining with anti-F4/80 for flow cytometry analysis. Cells were then acquired on FACS Celesta for 30 minutes after every 5 min and median fluorescent intensity of cypHer5E Alexa Fluor 647 on CTV^+^ F4/80^high^ macrophages was recorded. Acidification index was quantified as log fold change of median fluorescent intensity of cypHer5E on CTV^+^ F4/80^high^ macrophages at each indicated time over median fluorescent intensity of cypHer5E of F4/80^high^ macrophages at time 0 min.
- Confocal imaging of peritoneal macrophages: LPM were obtained from naïve WT or Tim4 KO animals and plated on ibidi chambers in IMDM for 6 hours. LPM were then pulsed with apoptotic ID8 ZsGreen^+^ for 15 minutes at 37°C, followed by three washes with PBS to remove non-bound thymocytes. After 15, 30 or 45 minutes of chase at 37°C, cells were fixed with 4% PFA for 5 minutes at room temperature. Blocking was performed with 5% mouse serum followed by staining with indicated antibodies. Nucleus was stained with Hoechst 33342 (Invitrogen) and mounting medium (ibidi) was added to preserve the samples until acquired. All the steps were performed in ibidi chambers. Confocal images were acquired with a LSM 880 META reverse microscope (Zeiss) with a 63×/1.4 NA plan oil objective. Image analyses were performed using Volocity 3D Image Analysis Software 5.5.1 (PerkinElmer) and Fiji (NIH). Curvature of actin phagocytic cup was calculated with the Kappa: a Fiji plugin for curvature analysis. To quantify actin surrounding cargo, within each cell (macrophage) we computed annular area of width 0.8 μm around cargo and measured actin integrated density followed by normalization with actin density per cell (macrophage). ‘Average thymocyte area’ was obtained from calculating area of n>20 uninternalized thymocytes. To quantify cargo degradation, scoring was done based on individual cargo area. “Intact”: Average thymocyte area ± 10%, “Partially digested”: Between 30-90%±10% of Average thymocyte area, “Completely digested”: <20%±10% of Average thymocyte area.
- Detection of OVA-MHC-I complexes on surface of peritoneal macrophages To detect specific MHC class-I OVA complexes peritoneal cells from tumor-bearing mice were incubated for 1⍰h at 37⍰°C with PE-coupled H2K^b^-SIINFEKL mAb (25-D1.16) (BioLegend). Cells were washed and stained with extracellular antibodies to identify LPM and SPM and analysed by flow cytometry. As negative control, LPM were isolated from control mice. As positive control, LPM and SPM isolated from control mice were stimulated for 2 hr with 50nM SIINFEKL and stained as above.
- Adoptive transfer of OT-I CD8 T cells and in vivo proliferation 1⍰×⍰10^6^ CD45.1^+^OTI CD8^+^ OVA-specific T cells were labeled with CFSE (5⍰µM, Biolegend) or CTV (5⍰µM, Thermo Fisher Scientific), and intraperitoneally injected into WT or Tim4 KO mice. After 16 hours, mice were injected with 2×10^5^ apoptotic ID8-OVA. For experiments with liposome-mediated antigen delivery, OVA-containing PC or PS liposomes amounting to equal concentrations of OVA (1-2 μM). Two days after tumor or liposomes challenge, mice were sacrificed and peritoneal ascites were collected, processed as described previously and proliferation was verified by flow cytometry. As a control, total ascites cells were re-stimulated ex-vivo with OVA class-I peptide SIINFEKL (2⍰µM).
- Activation of endogenous anti-tumor CD8^+^ T responses For analysis of the activation state of endogenous tumor-infiltrating CD8^+^ T cells in ID8^ZGO^tumor bearing mice, peritoneal cells were collected at day 14 post-tumor inoculation. Cells were washed with PBS and stimulated with SIINFEKL (2⍰µM) 37⍰°C for 4 h in the presence of Golgi Stop (BD Biosciences) to allow accumulation of intracellular cytokines. After viability and surface marker staining, cells were fixed and permeabilized using Cytofix/Cytoperm solution (BD Biosciences) following manufacturer’s instructions, and then stained with anti-IFNγ-PE (XMG1.2, Biolegend). Tim4 KO mice or WT counterparts were injected with 2 × 10^6^ ID8^ZGO^ cells and sacrificed 14 days post-tumor inoculation. The accumulation of endogenous anti-tumor CD8^+^ T cells in the peritoneal cavity was assessed by using Pro5® MHC H-2Kb Pentamers (Proimmune) following manufacturer’s instructions. Briefly, total peritoneal cells were stained with Pro5® MHC H-2KbPentamers for 45⍰min at 4⍰°C, washed and stained to identify CD8 using the gating strategy shown in Supplementary Fig. S1A.

### QUANTIFICATION AND STATISTICAL ANALYSIS

All statistical analyses were performed using GraphPad Prism 9 (GraphPad software Inc.). Data are presented as the mean ± SEM, unless otherwise indicated. The number of replicates for each experiment are mentioned in the respective figure legends. Statistical significance between two was evaluated using Student’s two-tailed t test (paired or unpaired, as applicable). For multiple comparisons, multiple t-test (Holm-Sidak correction), one-way ANOVA or two-way ANOVA followed by Tukey’s or Dunnett’s post-tests were performed as appropriate. For acidification and degradation curves, a simple linear regression model was applied and two groups were compared based on their slopes. P values < 0.05 were considered significant.

### KEY RESOURCES

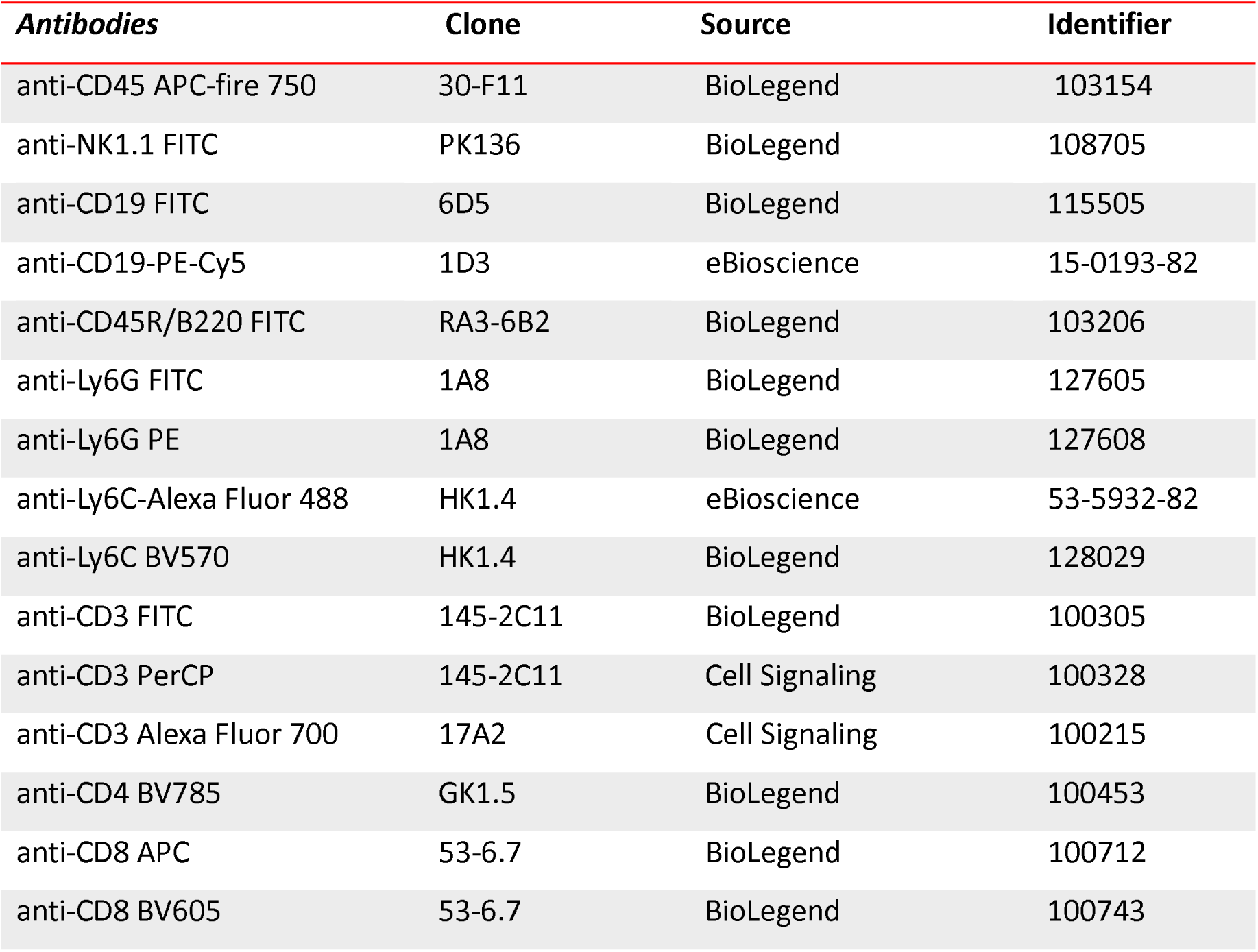
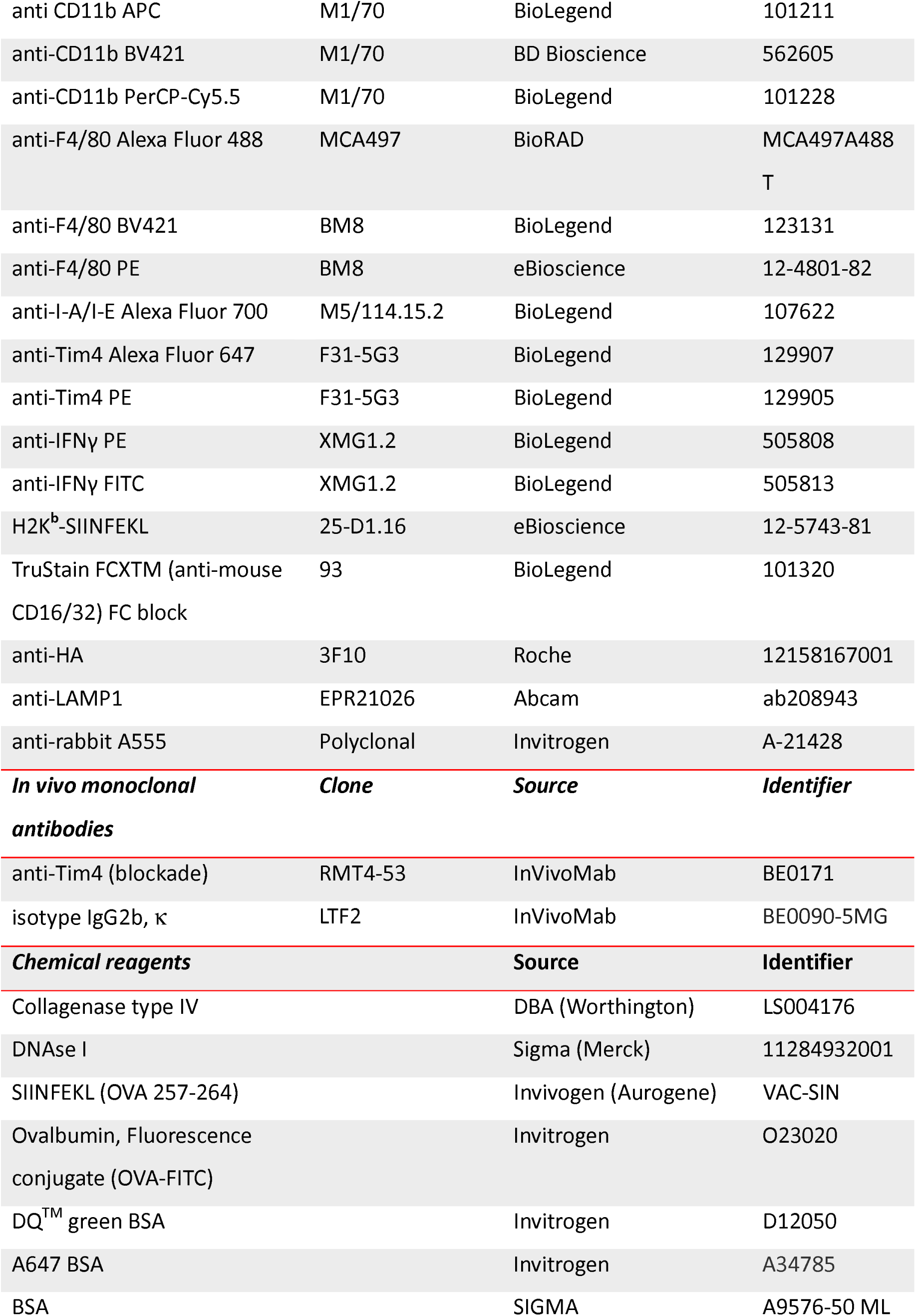
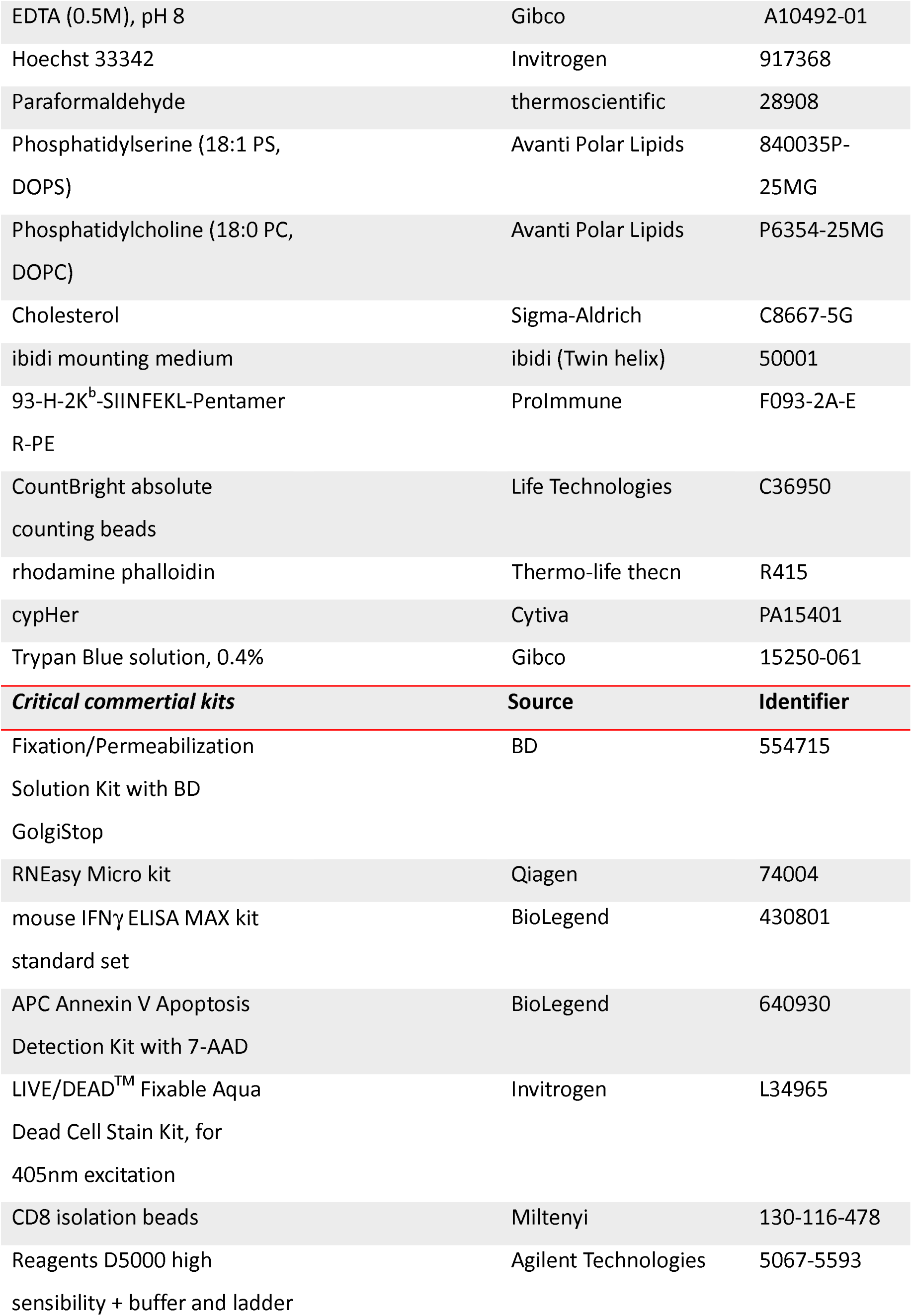
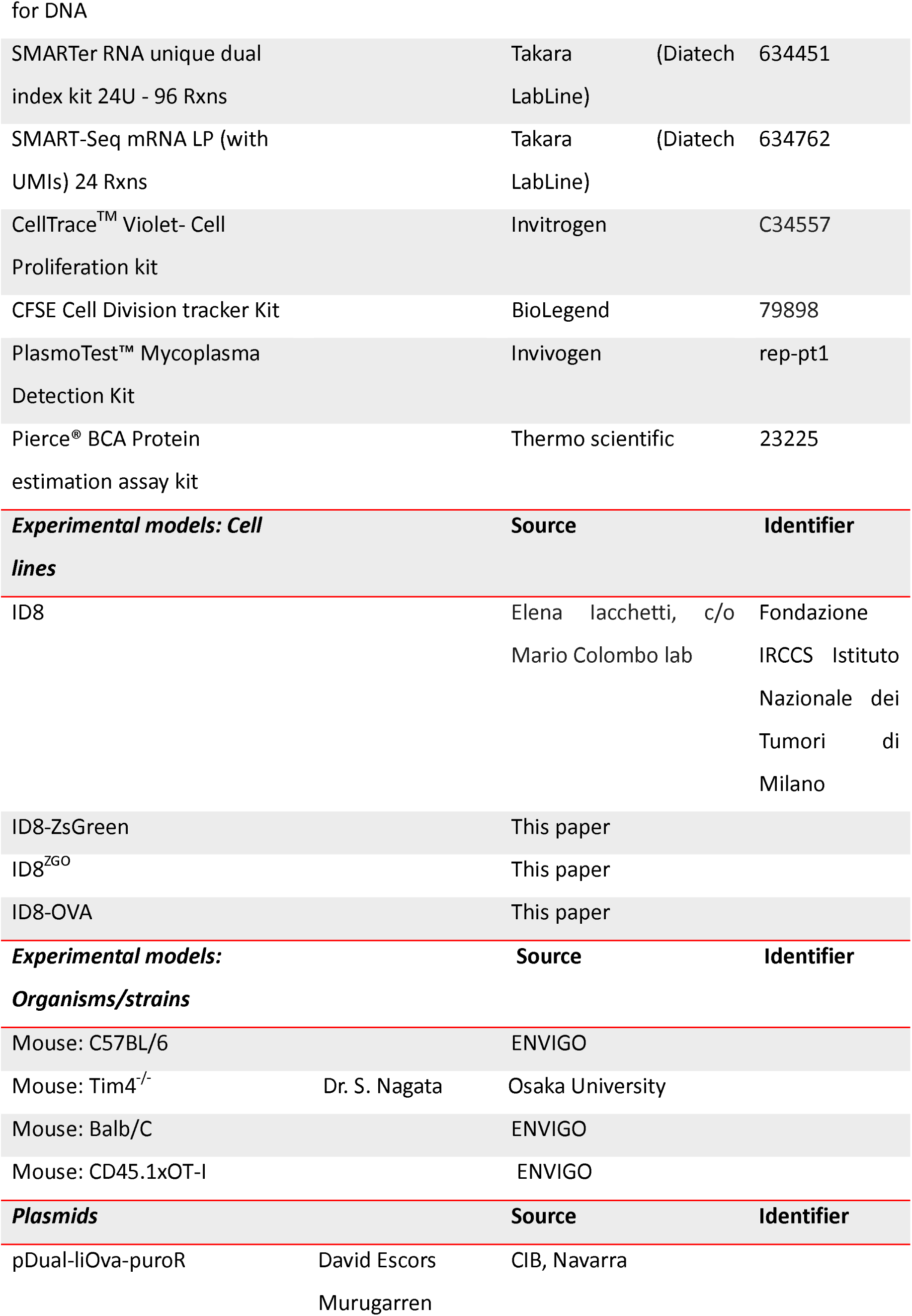
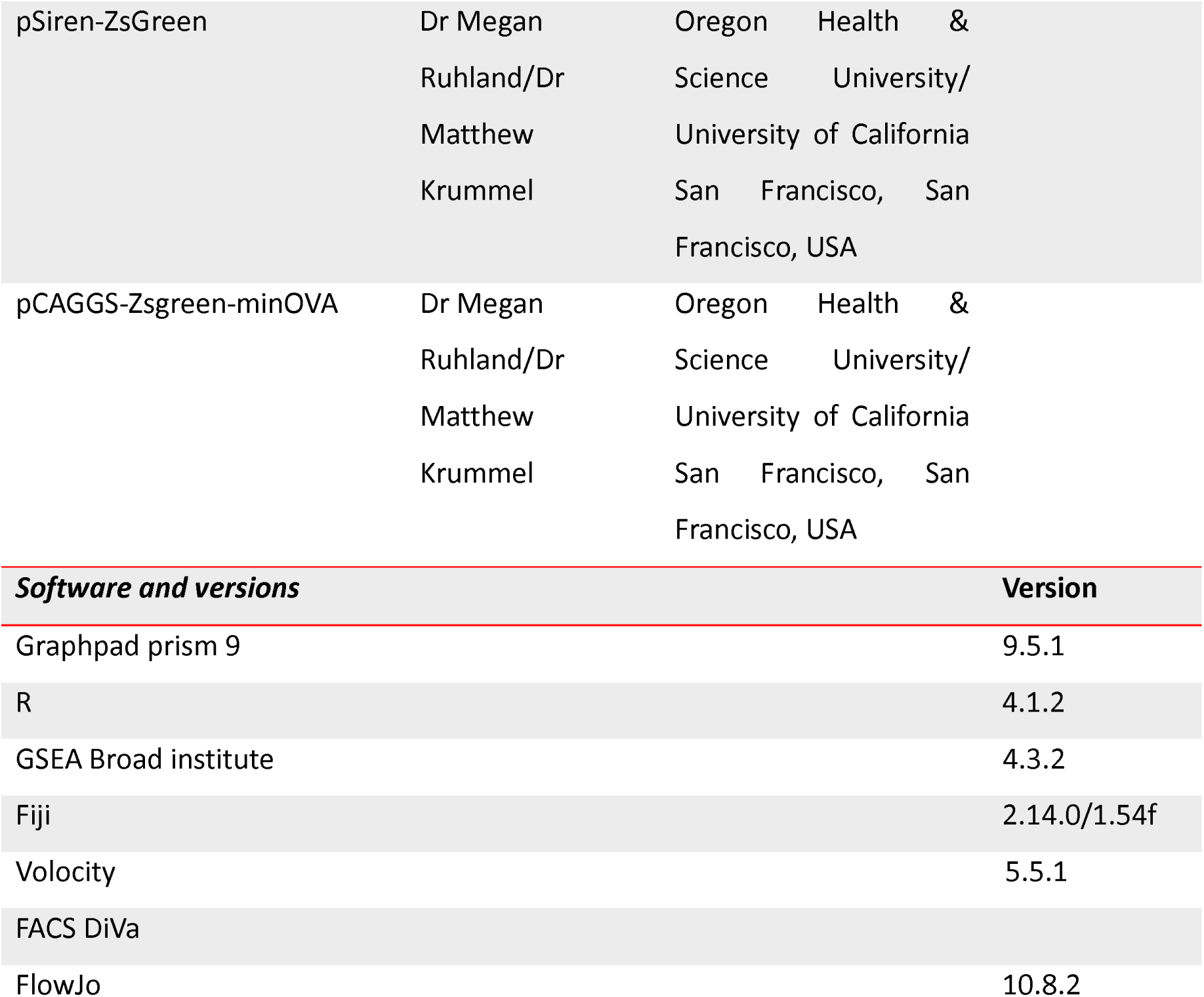

## Supplementary Tables

**Supplementary Table 1:** Pairwise significant DEGs Large Peritoneal macrophages

**Supplementary Table 2:** Pathway enrichment analyses of Large Peritoneal macrophages

