## Supplementary information for "Tim4 enables large peritoneal macrophages to cross-present tumor antigens at early stages of tumorigenesis"

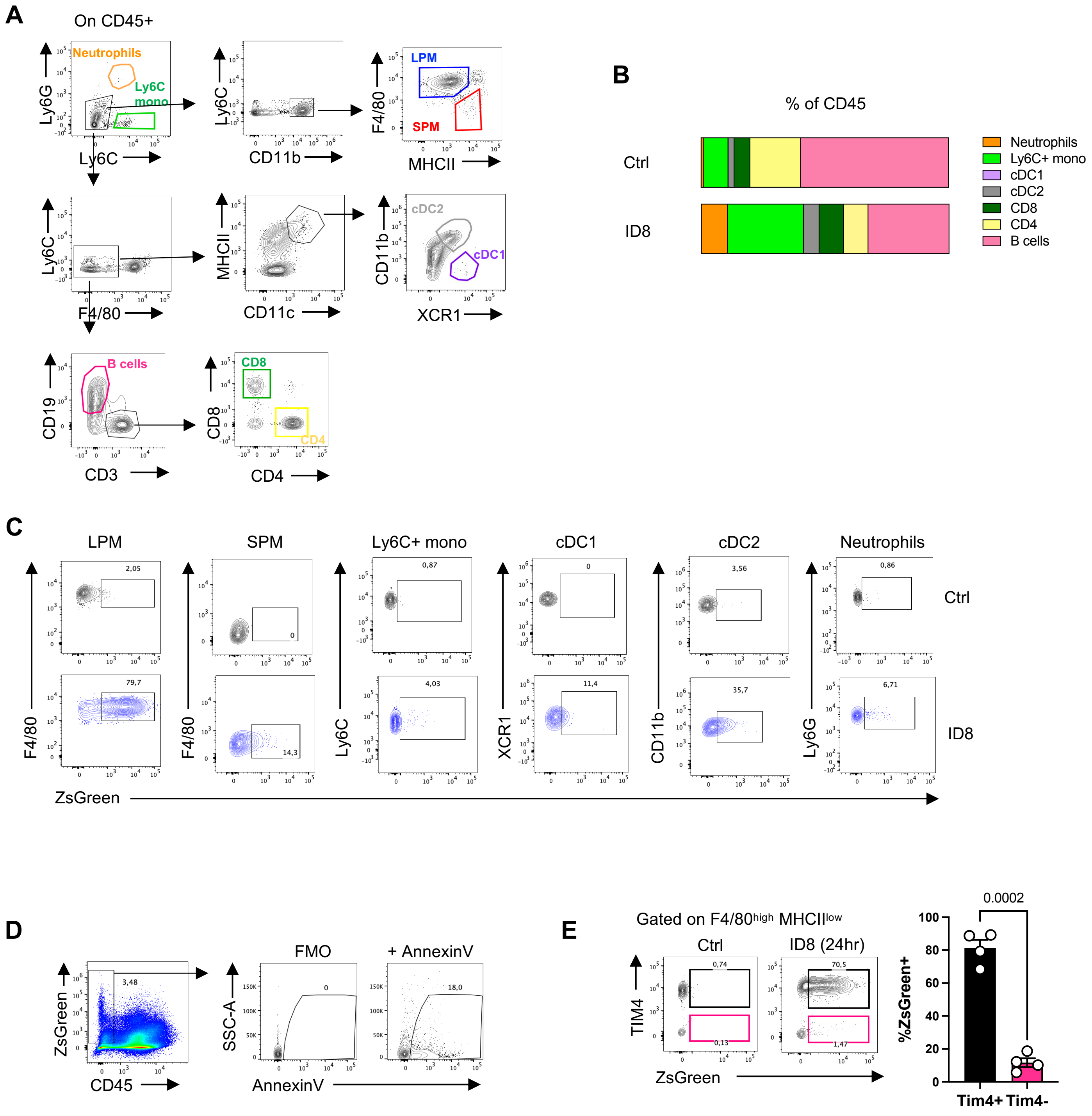


**Fig S1. Analysis of peritoneal macrophages in ID8 tumors.**

**Fig S1| Analysis of peritoneal macrophages in ID8 tumors.**

**A)** Gating strategy used to identify different immune cell subsets in the peritoneal cavity. **B)** Frequencies of indicated immune subsets in peritoneal cavity of resting and $ID8^{ZGO}$-challenged animals (day 15) plotted as percentage of total CD45. n=4. **C)** Dot plots showing uptake of $ID8^{ZGO}$ by the indicated phagocytic populations in the peritoneal cavity of naïve and tumor challenged mice. Representative of one of the 3 independent experiments with n=3-4 animals each. **D)** Phosphatidylserine expression on tumor cells ($CD45^-$ $ZsGreen^+$) isolated from the peritoneal cavity 15 days after $ID8^{ZGO}$ challenge by Annexin staining. **E)** Representative dot plots and quantifications of tumor cells uptake by $TIM4^+$ (black) and $TIM4^-$ (pink) LPMs 24 hr after tumor challenge. n=4, data represent mean ± SEM. Statistical analysis was conducted by paired t-test**.**

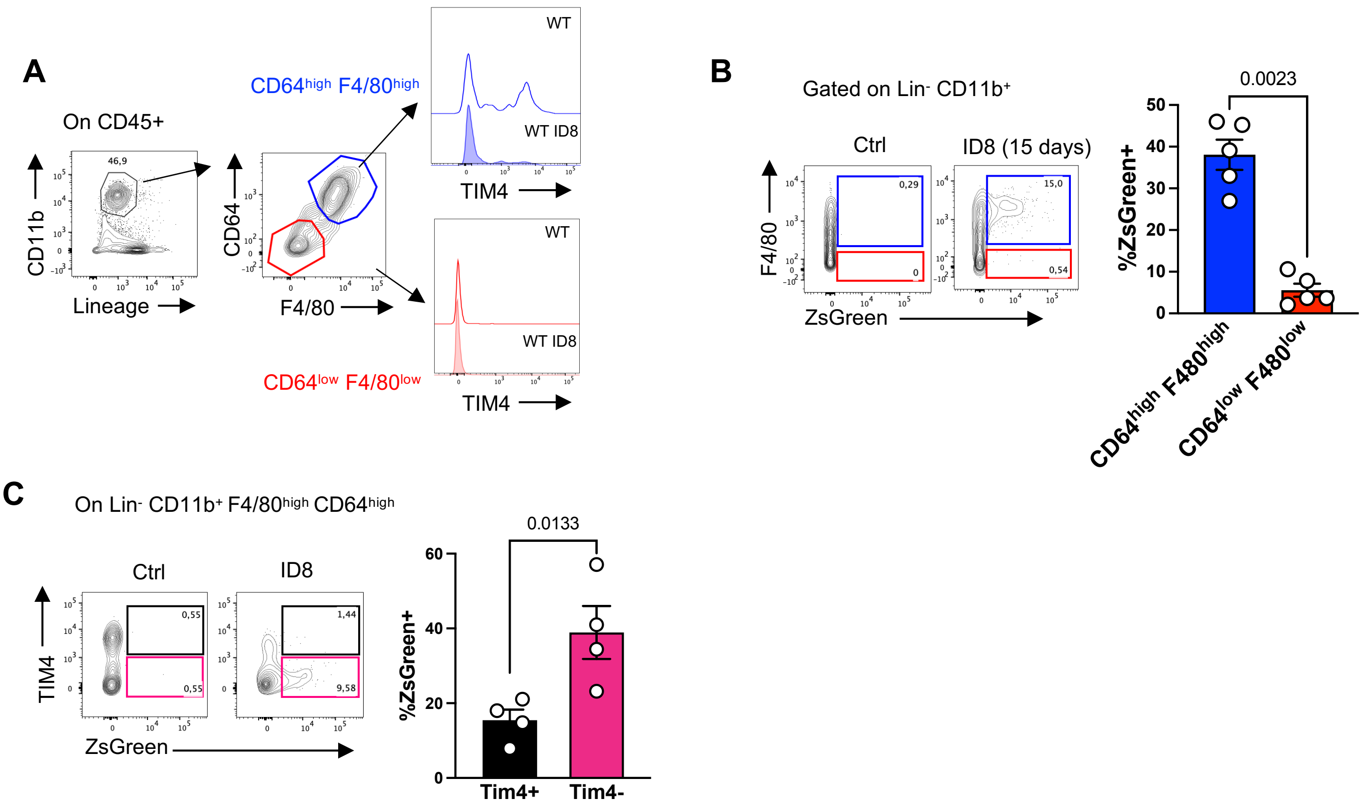


**Fig S2. Analysis of omental macrophages in ID8 tumors.**

**Fig S2| Analysis of omental macrophages in ID8 tumors.**

**A)** (left) Gating strategy to identify omental macrophage subsets ($CD64^{high}$ $F4/80^{high}$ & $CD64^{low}$ $F4/80^{low}$) and (right) histograms depicting TIM4 expression on respective subsets in naïve and $ID8^{ZGO}$ challenged mice (day 15). **B)** (left) Uptake of tumor cells by $F4/80^{high}CD64^{high}$ (blue) and $F4/80^{low}$ $CD64^{low}$(red) omentum macrophages (right) plotted as fraction of ZsGreen+ cells within each population. N=5, Data represent mean ± SEM. Statistical analysis was conducted by paired t-test. **C)** (left) Dot plots showing the uptake of tumor cells by TIM4+ (black) and TIM4- (pink) fractions of $F4/80^{high}$ $CD64^{high}$ omental macrophages and the relative quantification (right). n=4, Data represent mean ± SEM. Statistical analysis was conducted by paired t-test.

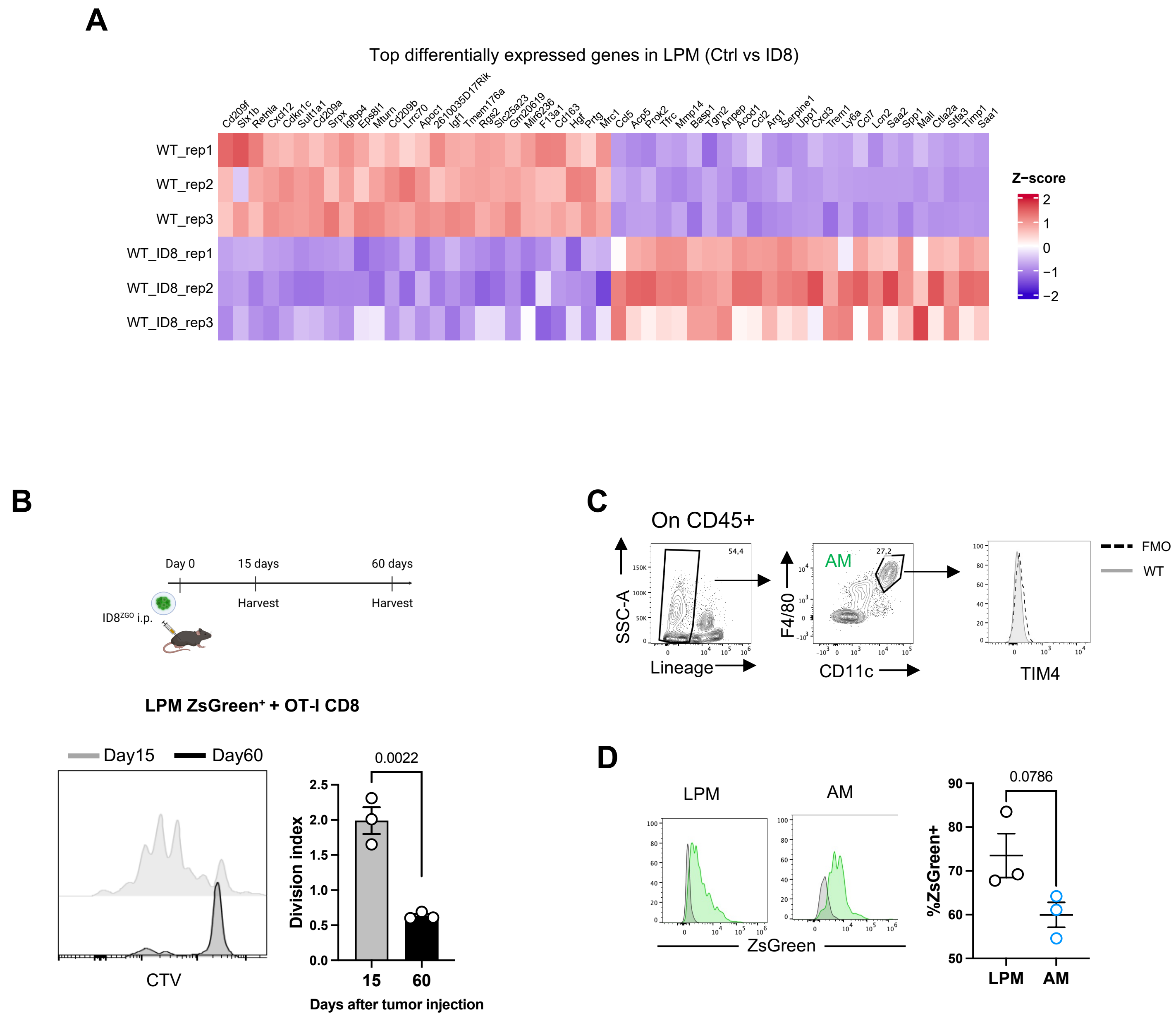


**Fig S3. Differential gene expression induced by phagocytosis of cancer cells, cross-presentation at late time points and in AM.**

**Fig S3| Differential gene expression induced by cancer cells uptake and cross-presentation at late time points and in AM. A)** Heat map showing z-scores of DEGs (top 30 and bottom 30) ($P < 0.05$, log2 (expression in WT ID8/expression in WT) > 1.5. **B)** Mice were injected with ID8$^{ZGO}$ intraperitoneally and ZsGreen+ LPM were isolated 15 or 60 days after tumor challenge. Sorted LPM ZsGreen+ were co-cultured with CTV labelled OT-I CD8 T cells and the fraction of proliferated cells was analyzed by flow and plotted as division index. n=3, Data represent mean ± SEM. Unpaired t-test. **C)** Gating strategy to identify Alveolar Macrophage subset (AM) from naïve mice and TIM4 expression. **D)** Cell-sorted LPM and AM were pulsed with ID8$^{ZGO}$ for one hour, washed and analyzed by flow cytometry. (left) Histograms represent ZsGreen fluorescence intensity in each population (green) overlaid with that of non-pulsed macrophages (gray). Efficiency of phagocytosis by each population (right) is plotted as fraction of ZsGreen$^{+}$ cells within each population. n=3, Data represent mean ± SEM. Statistical analysis was conducted by unpaired t-test.

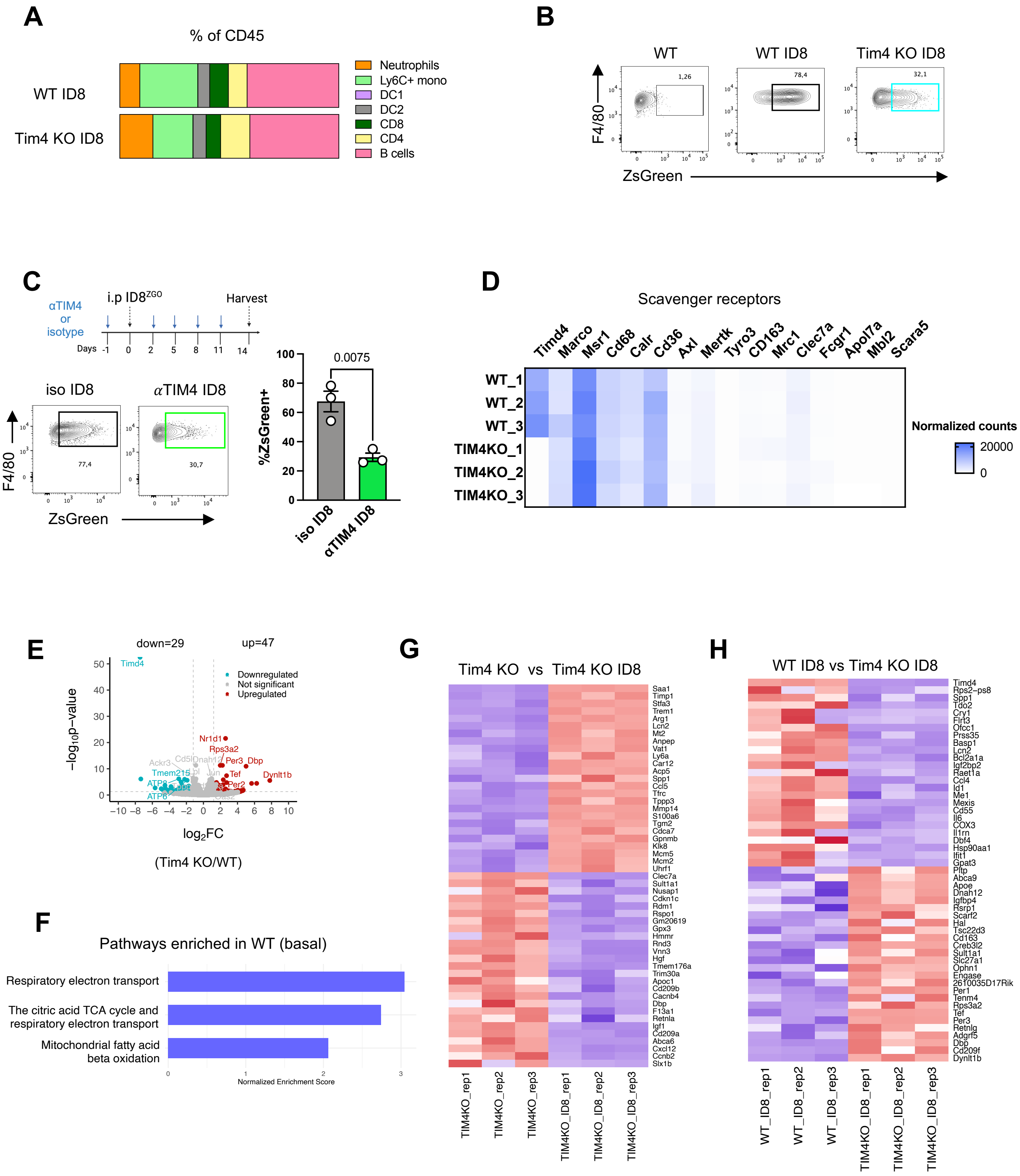


**Fig S4. Transcriptional remodeling in ZsGreen⁺ Tim4 KO LPMs**

**Fig S4| Transcriptional remodeling in ZsGreen$^{+}$ Tim4 KO LPMs**

**A)** Abundance of each of the indicated immune cell population in the peritoneal cavities of WT and Tim4 KO mice 15 days after challenge with ID8$^{ZGO}$, plotted as fraction of CD45+ cells. n=4. **B)** Dot plots representing uptake of ID8$^{ZGO}$ by LPM from WT and Tim4 KO mice 15 days after tumor challenge from **Fig 3C**. **C)** Analysis of phagocytic capacity of LPM under Tim4 blockade. (Upper panel) WT mice were injected with TIM4-blocking antibody one day before ID8$^{ZGO}$ peritoneal challenge and every 72 hours until day 15, on which peritoneal LPM were harvested. (Lower panel, left) Dotplots display capacity of LPM in to engulf ID8$^{ZGO}$ in presence or absence of Tim4 blockade analysed by flow cytometry and (lower panel, right) plotted as fraction of ZsGreen+ cells within LPM. n=3, data represent mean ± SEM from one of the two independent experiments. Statistical analysis was conducted by unpaired t-test. **D)** Normalised counts of indicated scavenger and phagocytic receptors from DEseq2 analysis of transcripts from WT or Tim4 KO LPM at basal level. Libraries were prepared from n=3 animals per group. **E)** Volcano plots displaying differentially expressed genes in LPM Tim4 KO as compared to WT at basal level, without tumor challenge. (Log2FoldChange threshold >1.5 or <-1.5, p value >0.05). **F)** Barplot of selected significantly enriched pathways in LPM from WT in comparison with Tim4 KO animals. **G-H)** Heatmaps representing z-scores of top 25 most differentially expressed genes in LPM from **G)** Tim4 KO naïve and tumor challenged (Tim4 KO ID8) and **H)** WT ID8 and TIM4 KO ID8 animals. Libraries prepared from n=3 mice per group.

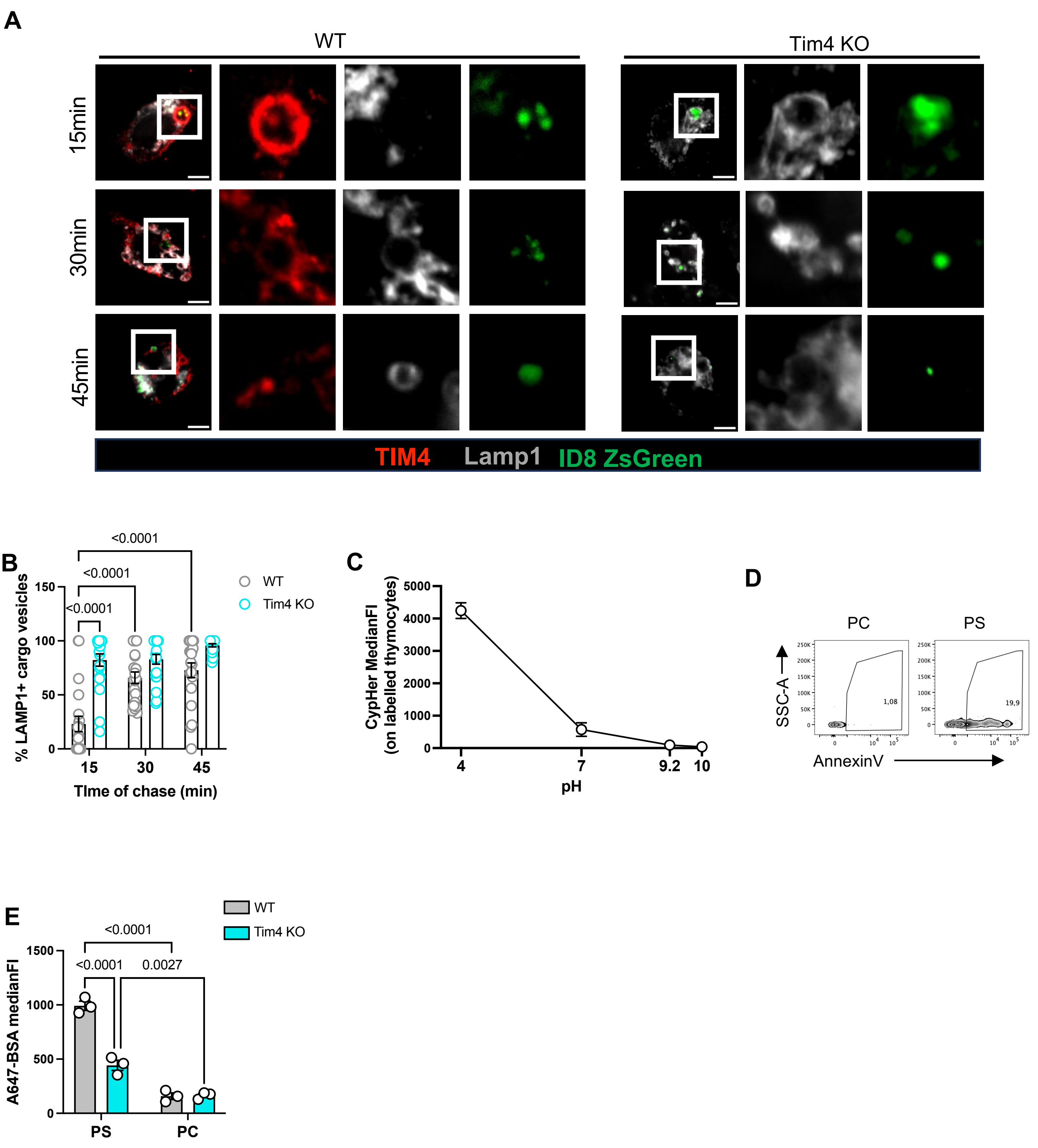


### Fig S5. Cancer cell debris are degraded faster in Tim4 null LPMs

**Fig S5| Cancer cell debris are degraded faster in Tim4 null LPMs**

**A)** LPM isolated from WT and Tim4 KO mice were pulsed for 15 minutes with apoptotic ID8 ZsGreen, chased for up to 45 minutes. Cells were fixed at 15, 30 or 45 minutes of chase time and later stained for Tim4 and Lamp1. Confocal images are representative of cells in each genotype at each time point. Scale bar=5μm. **B)** At indicated time points, fraction of tumor cell cargo within LAMP1 marked vesicles quantified in each cell cell. n>15 cells, data represent mean ± SEM from one of the two independent experiments. two-way ANOVA followed by Holm-Šídák's multiple comparisons test **C)** cypHer-labelled thymocytes were mixed with standard buffers of calibrated pH (4, 7, 10) or 0.1M sodium carbonate buffer (pH 9.2) and fluorescence intensity of cypHer was recorded by flow cytometry. Data are mean ± SEM. **D,E)** Characterization of DQ-BSA encapsulating liposomes. Lipo-PC and Lipo-PS were stained with Annexin-V and analyzed by flow cytometry. **E)** LPM isolated from WT and Tim4 KO mice were pulsed with either lipo-PC or lipo-PS encapsulating both Alexa Fluor 647-BSA and DQ-BSA for 15 minutes, followed by PBS-washes. Uptake of liposomes by LPM was analysed by flow cytometry after 20 minutes of chase time. The bars show the MFI of Alexa Fluor 647-BSA on F4/80$^{+}$ LPM. The data represents mean ± SEM from three independent experimental replicates. Statistical analysis was performed by two-way ANOVA followed by Tukey's multiple comparisons test.
